# The RNA-binding protein TRIM71 Promotes Supporting Cell Reprogramming and Hair Cell Regeneration by Activating *Sall4*

**DOI:** 10.64898/2026.10.05.756655

**Authors:** Lin Li, He Huang, Shixian Meng, Xingyu Liu, Yafan Wang, Ruifeng Qiao, Ying-Ying Xuan, Qin Hu, Xiao-yan Ma, Hui Ma, Shuhao Dong, Yinghui Wang, Yan Zhang, Bai Sun, Jun Li, Waldemar Kolanus, Angelika Doetzlhofer, Xiao-Jun Li

**Affiliations:** Frontier Institute of Science and Technology, Xi’an Jiaotong University, Xi’an 710049, China; Department of Otorhinolaryngology Head and Neck Surgery, The First Affiliated Hospital of Xi’an Jiaotong University, Xi’an 710061, China; The Solomon H. Snyder Department of Neuroscience, Johns Hopkins University School of Medicine, Baltimore, MD 21205, USA; School of Chemical Engineering and Technology, Tai Yuan University of Science and Technology, Shanxi 030024, China; Department of Laboratory Medicine, West China Biomedical Big Data Center, West China Hospital, Sichuan University, Chengdu, 610041, China; Department of Cell Biology and Genetics, Shanxi Medical University, Shanxi 030001, China; Department of Otolaryngology-Head & Neck Surgery, West China Hospital, Sichuan University, Chengdu 610041, China; Molecular Immunology and Cell Biology, Life & Medical Sciences Institute (LIMES), University of Bonn, Bonn, Germany

**Author notes:** **Correspondence addressed to:** Xiao-Jun Li, Ph.D., Frontier Institute of Science and Technology, Xi’an Jiaotong, University, Xi’an, 710061, China *, Angelika Doetzlhofer, Ph.D., Department of Neuroscience, Johns Hopkins University School of Medicine,. These authors contributed equally (Lin Li, He Huang).

## Abstract

Hair cell (HC) loss in the auditory organ (cochlea) is irreversible and represents the leading cause of deafness in humans. The absence of HC regeneration is primarily due to the limited plasticity of cochlear supporting cells (SCs). Using a novel doxycycline-inducible TRIM71 transgenic mouse model, this study demonstrates that overexpression of the RNA-binding protein TRIM71 promotes spontaneous cochlear HC regeneration in neonatal mice. Experiments in cochlear organoids indicate that TRIM71 reprograms SCs into progenitor-like cells and that this reprogramming is further enhanced by coactivation of follistatin (FST). FST augments TRIM71-induced increases in chromatin accessibility at progenitor and HC-specific gene loci, facilitating their transcriptional activation in SCs. Notably, FST also enhances TRIM71-induced reactivation of Sall4, which, like Trim71, is highly expressed in early otic progenitors. Using cochlear organoids and explants, we show that the early otic transcription factor SALL4 is required for TRIM71-induced HC formation. By contrast, SALL4 overexpression inhibits HC formation by maintaining reprogrammed SCs in an undifferentiated, proliferative state. These findings establish TRIM71 as an activator of HC regeneration in vivo, provide mechanistic insights into TRIM71’s role in SC reprogramming, and identify SALL4 as its essential effector.

**Significance Statement:** The mammalian cochlea lacks intrinsic capacity for hair cell (HC) regeneration, a deficiency that underlies the permanent nature of hearing loss. Using a novel inducible transgenic mouse model, we demonstrate that reactivation of Trim71, an otic progenitor gene, promotes HC regeneration in perinatal mice. TRIM71 induces an otic progenitor-like transcriptional program in supporting cells (SCs) through Sall4 activation, while concurrently initiating mesenchymal gene expression. This conflicting transcriptional state is resolved by co-activating follistatin, which suppresses mesenchymal conversion genes and increases chromatin accessibility at progenitor- and HC-specific gene loci. Collectively, our findings establish a mechanistic framework wherein TRIM71 functions as a key regulator of SC plasticity and identify the transcription factor SALL4 as a critical downstream effector in HC regeneration.

## Introduction

Hair cells (HCs) can be spontaneously regenerated from supporting cells (SCs) throughout the animal’s lifespan in non-mammalian vertebrates, such as birds and fish [reviewed in (1)]. In mammals, SCs in the auditory sensory organ (cochlea), lose the capacity to regenerate HCs after the onset of maturation, which in mice occurs at the end of the first postnatal week. The lack of cochlear HC regeneration in mammals leads to irreversible HC loss and constitutes a primary cause of permanent hearing loss in humans [reviewed in(2)]. Recent studies have shown that cochlear SCs, in response to HC loss, spontaneously regenerate a few HCs, particularly in the apical region of the cochlea(3, 4), and that this capacity can be enhanced by inhibiting Notch signaling and/or activating the WNT signaling pathway (5, 6). The HC-specific transcription factors ATOH1, POU4F3, and GFI1 are most often used for direct reprogramming of SCs into HCs (7–9), and have been successfully used to reprogram non-sensory cells into HCs in the mature cochlea (10). Moreover, combinatorial approaches that increase progenitor characteristics in SCs have been successful when combined with ATOH1 overexpression. Our recent study demonstrated that stimulation of JAG1/Notch signaling with JAG1-peptide enhances ATOH1-induced cochlear HC formation *in vivo* in 7-day-old mice (11). Similarly, transient activation of MYC and NOTCH in SCs boosts ATOH1-induced HC regeneration(12), and combined activation of MYC, NOTCH, WNT, and cAMP signaling efficiently reprogram SCs, leading to the regeneration of HC-like cells in the adult mouse cochlea (13). Despite these advances, regenerating functional HCs remains a major challenge, and no study to date has reported hearing recovery through HC regeneration in mammals. Thus, elucidating the regulatory mechanisms that govern cochlear SC plasticity is essential for uncovering the principles underlying HC regeneration.

We recently identified the RNA-binding protein TRIM71 as a critical regulator of cochlear HC development(14) and HC regeneration(15). TRIM71 is highly expressed during early animal development and is essential for neural tube closure and forebrain morphogenesis (16). TRIM71 haploinsufficiency leads to congenital hydrocephalus by altering the proliferation and differentiation of neural stem cells lining the embryonic cerebral ventricles (16), and a subset of patients present with sporadic malformations of the inner and middle ear, accompanied by hearing impairment (14). *Trim71* is highly expressed in otic progenitor cells (E.9.5-E11.5), and continues to be expressed at lower levels in inner ear HC and SC progenitors (prosensory cells) (E13.5), and after their differentiation, Trim71 expression is near undetectable (14). Loss of *Trim71* leads to premature withdrawal from the cell cycle and differentiation of cochlear prosensory cells, culminating in hearing loss. Mechanistically, *Trim71* deficiency upregulates *Inhba* and *Tgfbr2* expression and enhances p-SMAD2/3 signaling. Mouse genetic studies revealed that Activin A (Inhba) acts as a pro-differentiation signal (17), while TGFBR signaling limits prosensory cell proliferation (14).

In the developing cochlea, Activin and TGFBR signaling is counterbalanced by follistatin (FST), a secreted protein that antagonizes the biological activity of cytokines in the transforming growth factor-β (TGF-β) superfamily (18). Like *Trim71, Fst* is highly expressed during early otic development and is required to maintain cochlear prosensory cells in a proliferative and undifferentiated state (17). Furthermore, *Fst* is required for the maintenance of apical cochlear identity. Loss of *Fst* lead to a short cochlea without the apical domain and low-frequency-specific hearing loss (19). Similarly, *Trim71* knockout at E8.5 also impairs hearing in mice(14). However, the underlying mechanism about how *Fst* and *Trim71* regulate SC reprogramming and HC regeneration need to be further addressed.

Loss of *Trim71* attenuates the latent capacity of cochlear epithelial cells to form HCs in organoid culture, while its overexpression greatly enhances the ability of cochlear epithelial cells [SCs and greater epithelial ridge (GER) cells] to reenter the cell cycle, proliferate, and form HCs in cochlear organoid culture (15). Whether TRIM71 promotes HC regeneration *in vivo* is unknown. Furthermore, how TRIM71 reactivation enhances the regenerative plasticity of cochlear SCs at the cellular level, and the potential involvement of TGF-β signaling remain to be elucidated.

Here, by employing a novel doxycycline-inducible TRIM71 transgenic mouse, we show that reactivating TRIM71, together with FST, promotes HC regeneration in newborn mice. Characterization of TRIM71- and FST-induced transcriptomic and epigenetic changes reveals a critical role for FST in counterbalancing TRIM71-induced epithelial-to-mesenchymal transition, shows that TRIM71 and FST act synergistically to increase chromatin accessibility at progenitor- and HC-specific genes, and identifies SALL4 as a downstream effector of TRIM71. Knockdown of Sall4, which is co-expressed with Trim71 in the developing otocyst, attenuates HC formation in TRIM71 overexpressing cochlear organoid and explant cultures. Furthermore, we show that overexpression of SALL4 in cochlear organoids promotes cell proliferation but, by itself, is insufficient to enhance HC formation. In sum, our research identifies the TRIM71-SALL4 axis as a critical regulator of SC plasticity and HC regeneration.

## Results

### HC Formation in Cochlear Organoid Culture Established with *iTRIM71* transgenic mice

The organoid culture model overcomes the limited availability of SCs in vivo and has been widely used to investigate SC reprogramming and HC regeneration (20–22). In a recent study, we overexpressed TRIM71 in cochlear organoids and explants using a lentiviral and AAV vector, respectively (15). To further investigate the function of TRIM71 in HC regeneration, we generated doxycycline (Dox)-inducible *TRIM71* overexpression mice using CRISPR/Cas9. We inserted a fusion protein expression cassette driven by a TRE promoter into the TIGRE safe-harbor site of the mouse genome. This cassette contains a 3×HA tag, the human TRIM71 coding sequence (CDS), a P2A self-cleaving peptide, and the tdTomato red fluorescent reporter gene (*SI Appendix*, Fig. 1A). The red fluorescence of the tdTomato protein was used to monitor TRIM71 expression. Expression of the reverse tetracycline transactivator transgene (*M2rtTA*) together with the *TRIM71* transgene (referred to as *iTRIM71*) allows us to overexpress TRIM71 by administering doxycycline (Dox) (*SI Appendix*, Fig. S1*B*).

**Fig. 1.**
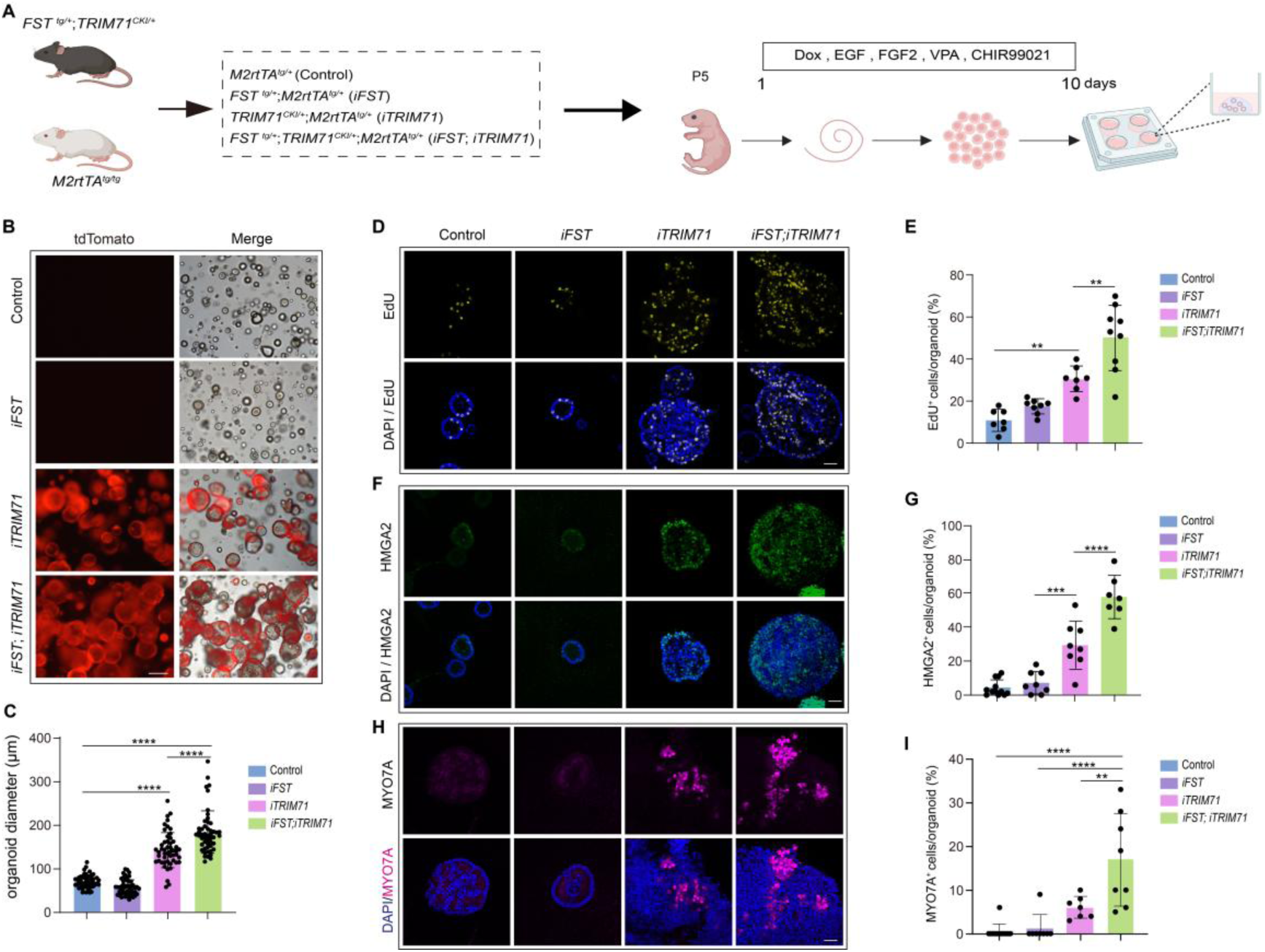
Concurrent FST and TRIM71 overexpression promotes HC formation in cochlear organoid cultures at stage P5. (*A*) Experimental scheme. organoid culture was established with cochlear epithelial cells from P5 control, *iFST*, *iTRIM71*, and *iFST;iTRIM71* transgenic mice. Created in BioRender. (*B*) Low-power bright-field and read fluorescent (tdTomato) images of organoid cultures. Scale bars=200 μm. (*C*) The organoid diameter in (*B*). (*D*) Cell proliferation in control, FST-, TRIM71-, and FST+TRIM71-overexpressing organoids. A single EdU pulse was given at 10 days of expansion, and EdU incorporation (yellow) was analyzed 0.5 hour later. DAPI (blue) staining marked cell nuclei. Scale bars=40 μm. (*E*) Percentage of EdU^+^ cells in (*D)* (n=7-9, two independent experiments). (*F*) Organoids were immunostained for HMGA2 (green). Scale bars=40 μm. (*G*) Percentage of HMGA2^+^ cells per organoid in (*F*). (*H*) High-power images of HC marker MYO7A (magenta). Scale bars=40 μm. (*I*) Percentage of MYO7A^+^ cells per organoid in (*H*). One-way ANOVA and Tukey’s multiple comparison test were performed for *C*, *E*, *G* and *I*. *\*P<0.05, **P<0.01, ***P<0.001, ****P<0.0001*.

To validate the new *iTRIM71* mouse model, we conducted TRIM71 overexpression experiments in cochlear organoid cultures. Cochlear epithelial cells, composed of SCs and GER cells, were isolated from stage P5 control (*Atoh1-nGFP*) and *iTRIM71-*transgenic (*iTRIM71;Atoh1-nGFP*) mice (SI Appendix, Fig. S1*C*). To induce cell cycle reentry and sustain cell proliferation, cochlear epithelial cells were placed in a Matrigel and cultured in a proliferation medium containing Dox, as well as epidermal growth factor (EGF), fibroblast growth factor 2 (FGF2), a TGF-βRI inhibitor (616452), the HDAC inhibitor valproic acid (VPA), and a GSK3 inhibitor (CHIR99021), to boost WNT signaling (*SI Appendix*, Fig. S1*C*). Validating the newly developed *iTRIM71* mouse model, we found that, similar to viral overexpression, transgenic overexpression of *TRIM71* induced robust HC formation, with the percentage of Atoh1-nGFP⁺ organoids reaching 40%, whereas in the control group only 4% Atoh1-nGFP^+^ organoids were detected (SI Appendix, Fig. S1D and *G*). On day 10 of expansion, TRIM71 overexpression significantly increased the diameter and colony-forming number of organoids compared with the control (SI Appendix, Fig. S1E and *F*).

### Concurrent FST and TRIM71 Overexpression Enhances SC Proliferation and HC Formation in Cochlear Organoid Culture

The RNA-binding protein LIN28B, a well-known antagonist of the *let-7* family of microRNAs, is a key regulator of cellular plasticity and pluripotency [reviewed in (23)]. In the cochlea, LIN28B reactivation boosts SC plasticity and HC regeneration by reprogramming SCs into progenitor-like cells, an effect further enhanced by co-activation of the Activin antagonist FST (24). To investigate whether FST+TRIM71 coactivation has a similar positive effect as FST+LIN28B overexpression, we generated control (*M2rtTA^tg/+^*), *iFST* (*FST^tg/+^;M2rtTA^tg/+^*), *iTRIM71* (*TRIM71^CKI/+^;M2rtTA^tg/+^*) and *iFST;iTRIM71* (*FST^tg/+^;TRIM71^CKI/+^;M2rtTA^tg/+^*) mice (Fig. 1*A*), harvested their cochlear epithelial cells at postnatal day 5 (P5) and established organoid cultures. To avoid interference, we omitted the TGFBR1 inhibitor 616452 from the culture medium. After 10 days of expansion, we found that the average diameter of organoids was significantly greater in TRIM71-overexpressing cultures than in control cultures. Notably, the average diameter of FST+TRIM71-overexpressing organoids was further increased compared with that of TRIM71-overexpressing organoids (Fig. 1 *B* and *C*). This was consistent with the increase in cell proliferation observed in organoids overexpressing FST+TRIM71 compared with TRIM71-only (Fig. 1 *D* and *E*).

Our recent studies revealed that overexpression of LIN28B or TRIM71 in cochlear organoid cultures reprograms SCs into progenitor-like cells and enhances HC formation. This process is driven in part by the activation of HMGA2 expression, a chromatin-binding protein (24, 25). In the developing cochlea, *Hmga2* is highly expressed in prosensory cells and is rapidly downregulated during differentiation (26). Consistent with FST acting as a cofactor in TRIM71-mediated SC reprogramming, we found that the percentage of HMGA2⁺ cells in FST+TRIM71-overexpressing organoids (57%) was nearly 2-fold higher than in TRIM71 alone (30%) (Fig. 1 *F* and *G*). Furthermore, consistent with FST co-expression enhancing TRIM71-induced HC formation, we found that the percentage of MYO7A⁺ cells was significantly higher in FST+TRIM71-overexpressing organoids (17%) than in TRIM71-expressing organoids (6%). No MYO7A⁺ <u>HC</u>s were observed in control or FST-overexpressing organoids (Fig. 1 *H* and *I*).

### Concurrent FST and TRIM71 Overexpression Enhances SC Proliferation and HC Formation in Cochlear Explant Culture

Our previous study showed that AAV-mediated overexpression of TRIM71 enhanced HC formation in P2 cochlear explants in response to inhibition of Notch signaling (15). To validate this result and determine whether TRIM71 overexpression at later stages has a similar positive effect on HC formation, we established cochlear explant cultures from control and *iTRIM71* mice. The culture media contained Dox to induce TRIM71 transgenic expression and LY411575, a small-molecule Notch inhibitor, to trigger HC formation (Fig. 2*A*). After 4 days of culture, we observed SOX2^+^tdTomato^+^ cells in the SC layer of the TRIM71-overexpressing cochlea, indicating successful induction of TRIM71 in SCs (SI Appendix, Fig. S2). The number of MYO7A^+^SOX2^+^ cells in the middle and base of P2 TRIM71-overexpressing cochlear explants was more than 3-fold higher than in littermate controls (Fig. 2 *B-D*). However, TRIM71 overexpression failed to enhance HC formation in response to Notch inhibition in cochlear explants isolated from P5 mice (Fig. 2 *E-K*).

**Fig. 2.**
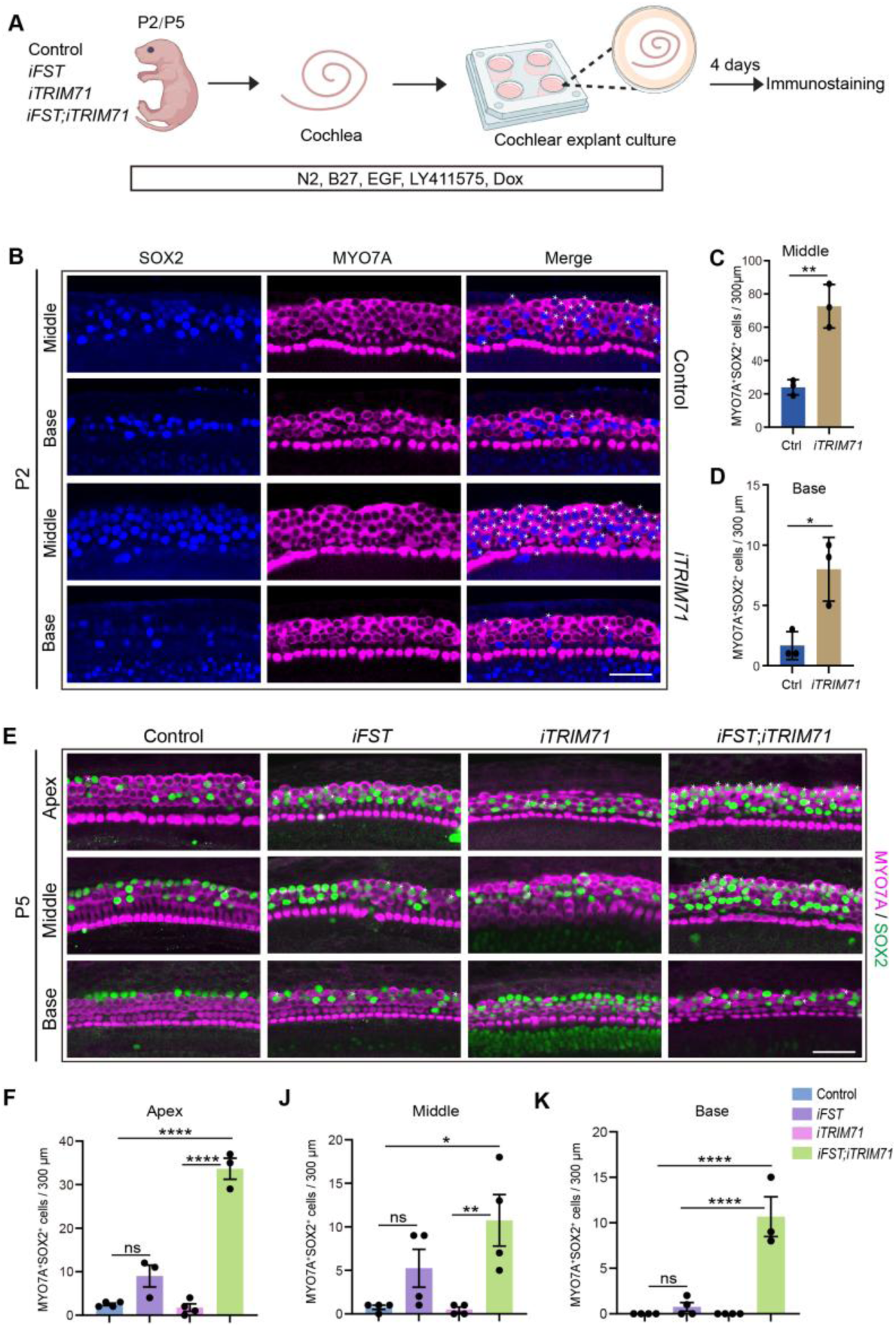
Coactivation of FST and TRIM71 promotes HC formation in P5 cochlear explant culture. (*A*) Experimental scheme. Cochlear sensory tissue from control, *iFST*, *iTRIM71,* and *iFST*;*iTRIM71* transgenic mice was isolated at stage P2 /P5 and cultured in the presence of EGF (5 ng/mL), LY411575 (5 μM), and dox (0.5 μg/mL) for 4 days. Created in BioRender. (*B*) Confocal images of the HC layer of control and TRIM71 (*iTRIM71*) overexpressing cochlear explants. Shown are representative images of the sensory epithelia at the cochlear mid and base. MYO7A immunostaining marks both existing and newly formed HCs. Co-labeling of MYO7A and SOX2 marks newly formed HCs (white asterisks). Scale bars=40 μm (*C-D*). Quantification of MYO7A^+^SOX2^+^ cells in (*B*) n=3 animals per group, two independent experiments. (*E*) Confocal images of the HC layer in control, FST (*iFST*), TRIM71 (*iTRIM71*), and FST+TRIM71(*iFST*;*iTRIM71*) overexpressing cochlear explants. Co-labeling of MYO7A and SOX2 marks newly formed HCs (white asterisks). Scale bars=40 μm (*F-K*). Quantification of new HCs (MYO7A^+^ SOX2^+^) formed in the cochlear apex, middle, and base. Individual data points in *F-K* represent the number of animals analyzed per group. Two-tailed, unpaired Student’s t test was used to calculate P values for *C-D*. One-way ANOVA and Tukey’s multiple comparison test were performed for *F-K*. *\*P<0.05*, *\*\*P<0.01*, *\*\*\*\*P<0.0001*. n.s., not significant.

To investigate whether coactivation of FST and TRIM71 allows for new HC formation in response to Notch inhibition at P5, we generated cochlear explant cultures from P5 control, *iFST*, *iTRIM71,* and *iFST;iTRIM71* transgenic mice (Fig. 2 *E-K*). No MYO7A⁺SOX2⁺ cells were detected in TRIM71-overexpressing and control cochlear explants (Fig. 2 *E* to *K*). There were a few newly formed HCs (MYO7A⁺SOX2⁺) cells in the apical, mid, and basal portions of cochlear tissue explants following overexpression of FST, which was not significantly different from control explants (Fig.2 *E* to *K*). However, large numbers of MYO7A⁺SOX2⁺ double-positive cells were detected across the entire length (from apex to base) of P5 cochlear explants following co-activation of FST and TRIM71 transgenes (Fig.2 *E* to *K*). In summary, we propose that TRIM71 alone is insufficient to enhance HC formation in P5 cochlear explant culture, whereas coactivation of FST can greatly enhance HC formation in response to TRIM71 overexpression.

### TRIM71 Reactivation Promotes Spontaneous Cochlear HC Regeneration in Perinatal Mice

Previous studies have shown that the loss of HCs in early postnatal mice triggers a regenerative response (3, 4). Such spontaneous HC regeneration occurs only at perinatal stages and is limited to the cochlear apex. To investigate whether TRIM71 is involved in this regenerative process, we first assessed whether *Trim71* expression is induced by HC loss/damage. We established cochlear explant culture from P5 mice, and HCs were killed/damaged by exposing the explants to ototoxic drug gentamicin for 12 hours. Cochlear epithelia were then harvested at 12 and 24 hours after the initiation of gentamicin treatment (SI Appendix, Fig. S3*A*). We found that 12 hours post-HC injury, *Trim71* was significantly increased in cochlear epithelial cells, accompanied by increased *Hmga2* expression (SI Appendix, Fig. S3*B*). Trim71 and Hmga2 expression declined between 12 to 24 hours post-HC injury but remained above baseline levels. These findings suggest that the transient activation of *Trim71* in SCs after damage serves as a key signal for spontaneous HC regeneration.

To further investigate the role of TRIM71 in spontaneous HC regeneration *in vivo*, we generated *Pou4f3^DTR/+^* (*DTR*) mice carrying the *iTRIM71* transgene and their control littermates. The inclusion of the DTR transgene enables selective ablation of HCs by administering diphtheria toxin (DT) (27). We administered Dox at P1 to induce the expression of the TRIM71 transgene, and DT was administered at stage P1 (Fig. 3*A*). As expected, there were very few, if any, regenerated HCs (MYO7A^+^SOX2^+^) in control and TRIM71-overexpressing cochleae without HC damage (Fig. 3 *B* and *C*). However, consistent with previous reports, HC damage triggered the generation of a few MYO7A^+^SOX2^+^ cells in the cochlear apex of *Pou4f3^DTR/+^* transgenic control animals (DTR). Furthermore, the number of MYO7A^+^SOX2^+^ cells in the cochlear apex, mid, and base was significantly higher in *iTRIM71;DTR* animals than in *DTR* control animals (Fig. 3 *B* and *C*).

**Fig.3.**
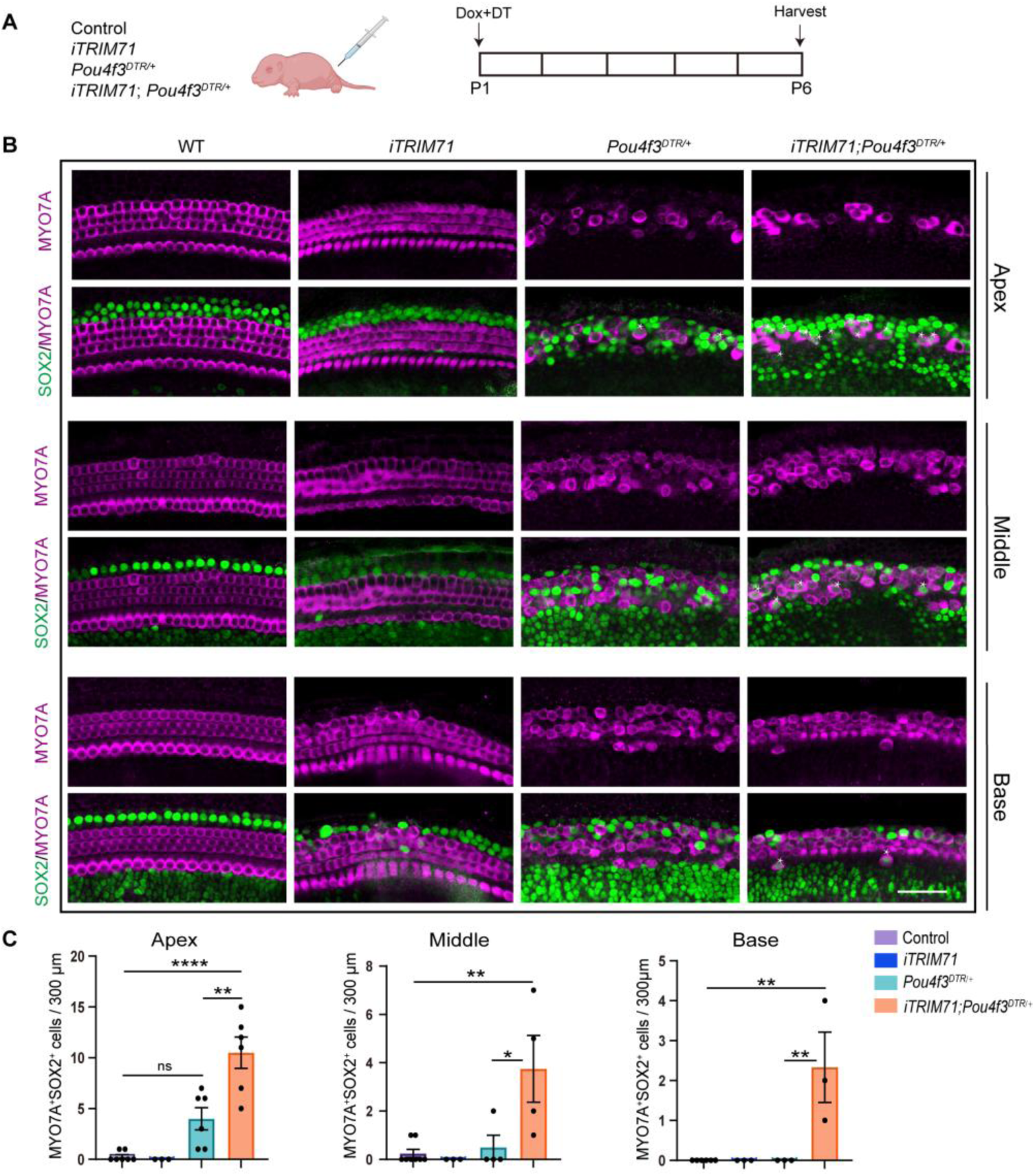
TRIM71 overexpression enhances cochlear HC regeneration in neonatal mice. (*A*) Experimental scheme. To induce TRIM71 expression in neonatal mice, mice were injected with Dox at P1, and cochlear tissue was harvested at P6. To ablate HCs, mice were injected with DT (6.25 ng/g) at stage P1. Created in BioRender. (*B*) High-power single-plane confocal images of cochlear sensory epithelia stained for MYO7A and SOX2. Scale bars=40 μm. (*C*) Quantification of newly formed HCs in the apex, middle, and base of control and TRIM71-overexpressing mice with or without HC damage. Individual data points represent the number of animals analyzed per group. One-way ANOVA and Tukey’s multiple comparison test were performed for *C*. *\*P*<0.05, *\*\*P*<0.01, and *\*\*\*\*P*<0.0001. n.s., not significant.

Mounting evidence suggests that in mice, cochlear SC plasticity rapidly declines in the first postnatal week (28, 29). To determine whether TRIM71 overexpression at later stages boosts spontaneous HC regeneration, we conducted experiments using stage P7 *iTRIM71;DTR* or *iFST;iTRIM71;DTR* mice. We found HC damage at P7 failed to trigger spontaneous HC regeneration in the TRIM71*-*overexpressing cochlea, with few sporadic new HCs in the apical region of the FST+TRIM71-overexpressing cochlea (*SI Appendix*, Fig. S4). In summary, these data indicate that TRIM71 overexpression enhances spontaneous HC regeneration in mice, but this positive effect is confined to early postnatal stages.

### FST and TRIM71 Reprogram SCs into Progenitor-like Cells Primed for HC Fate Induction

To identify how FST and TRIM71 coactivation boosts regenerative competence, we analyzed the transcriptomes of P5 control, TRIM71-, and FST+TRIM71– overexpressing organoids after 10 days of expansion using single-cell RNA sequencing (scRNA-seq). Libraries were prepared using the 10×Genomics platform. After stringent quality control, 19,050, 16,969, and 22,420 cells from control, TRIM71-overexpressing, and FST+TRIM71-overexpressing organoids, respectively, were harvested for scRNA-seq experiments. We identified 11 distinct cell clusters using Surat (v5.1.0) (Fig. 4*A*). Cell cluster 10 was identified as HCs based on the expression of well-known markers, including *Gfi1*, *Pou4f3* and *Tmem255b* (Fig. 4*B*). By comparison with previous inner ear organoid scRNA-seq data sets (30), cell cluster 0 was identified as greater epithelial ridge (GER)-like cells, characterized by expression of *Tecta*, *Sparcl1*, *Slc24a3*, and *Il33* (Fig. 4*B*). Cell clusters 1, 3, 5, and 8 showed expression patterns consistent with SC lineages but could not be assigned to specific SC subtypes; these clusters were thus classified as SC-like cells (Fig. 4*B*). Cell clusters 2, 4, 6, and 7 were classified as proliferative cells based on their high expression of mitotic marker genes, including Top2a and Ccna2. To characterize the proliferative state of cells across these clusters, we analyzed cell-cycle scoring using the Seurat CellCycleScoring function with canonical cell-cycle marker genes (31). The analysis revealed distinct proliferative patterns across clusters (Fig. 4*C*). Cluster 2 was highly enriched for mitotic cells (96.5% in G2/M phase), while cluster 4 was enriched for actively replicating cells (82.0% in S phase). Clusters 6 and 7 showed transitional profiles with mixed S and G2/M populations, suggesting active progression through late-cell-cycle phases. Control organoids were predominantly composed of GER-like cells (cluster 0) (Fig. 4 *A* and *C*) (SI Appendix, Fig. S5). Upon TRIM71 overexpression, the proportion of cells in SC-type clusters (1, 3, 5, 8) and clusters with proliferative cells (2, 4, 6) was increased, while the proportion of cells in clusters 0 and 7 was decreased, a pattern resembling that observed following coactivation of FST and TRIM71 (Fig. 4 *A* and *C*) (SI Appendix, Fig. S5). Furthermore, newly formed HCs (cluster 10) were found only in TRIM71-overexpressing and TRIM71+FST-overexpressing organoids (SI Appendix, Fig. S5).

**Fig.4.**
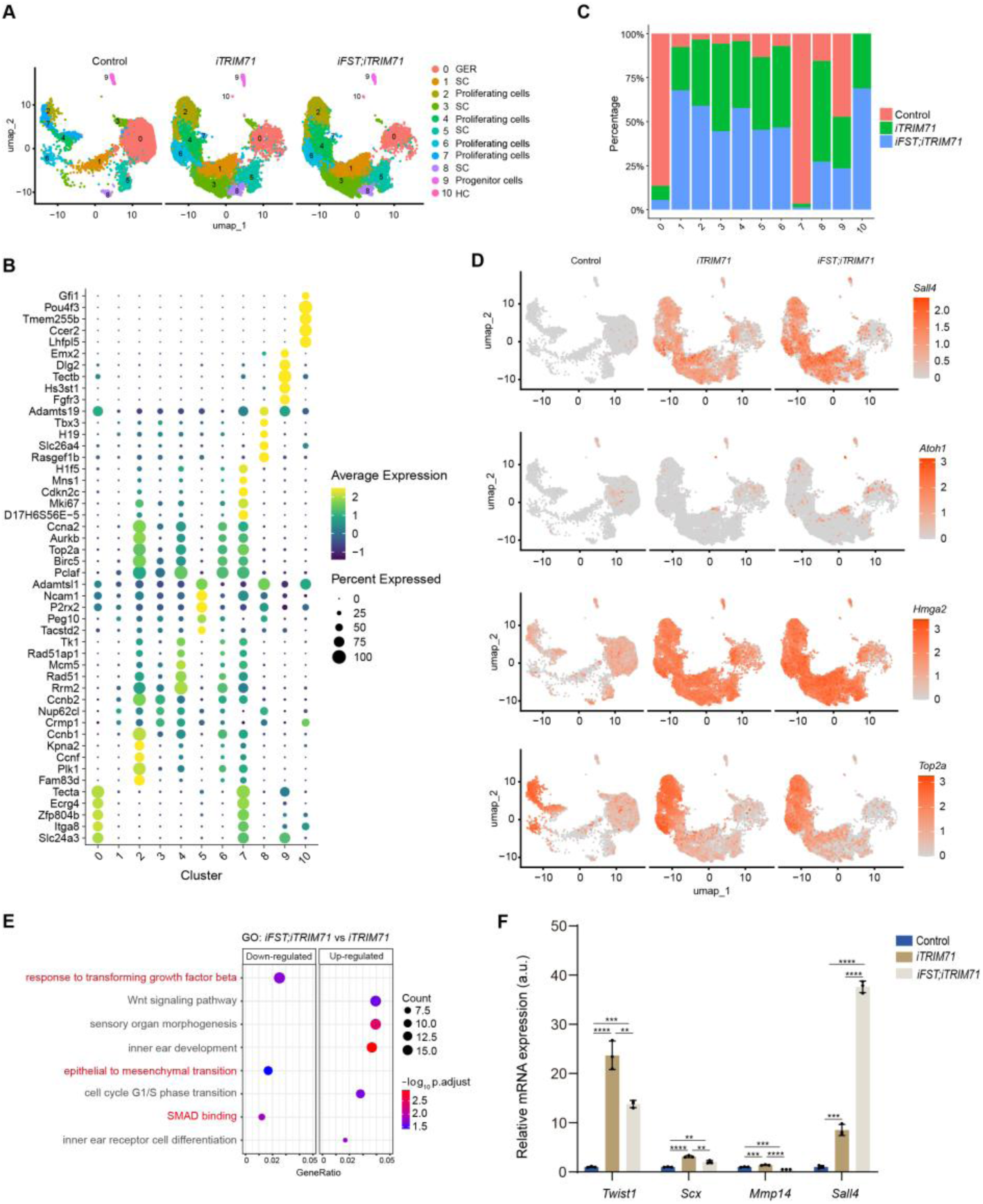
TRIM71 activates progenitor gene expression in cochlear organoid culture. (*A*) Uniform Manifold Approximation and Projection (UMAP) representing the shared cell clusters in the control (left), TRIM71-(middle), and FST+TRIM71-overexpressing organoids (right). (*B*) The top five differentially expressed genes for each cluster are displayed, with dot size representing expression percentage and color indicating expression level. (*C*) Bar plot showing the percentage distribution of each sample across identified clusters. (*D*) Feature plots for *Sall4, Atoh1*, *Hmga2* and *Top2a*. (E) GO enrichment of genes that are down-regulated and up-regulated in FST+TRIM71-overexpressing organoids compared with TRIM71-overexpressing samples. (F) RT-qPCR analysis of the expression of *Sall4* and mesenchymal genes (*Twist1*, *Scx*, *Mmp14*) in control, TRIM71, and FST+TRIM71-overexpressing organoids. One-way ANOVA and Tukey’s multiple comparison test were used to calculate P values. *\*P<0.05*, *\*\*P<0.01*, *\*\*\*P<0.001* and *\*\*\*\*P<0.0001*.

We found that progenitor genes such as *Sall4*(32) *and Hmga2*(24, 33), immature HC-specific gene *Atoh1* (34, 35), and cell-proliferating marker gene *Top2a* (36) were activated in TRIM71-overexpressing cells compared with control, and this effect was further enhanced in FST+TRIM71-overexpressing cells (Fig. 4*D*). The transcription factor SALL4 plays a central role in maintaining stem cell self-renewal (32) and has been reported to be involved in limb regeneration and wound healing in lower vertebrates(37). The remaining three members of the SALL transcription factor family, *Sall1*, *Sall2*, and *Sall3,* were not activated following TRIM71 overexpression compared with control (*SI Appendix*, Fig. S6). We analyzed published scRNA-seq data sets (38) and found that *Trim71* and *Sall4* are highly expressed in the otic vesicle cells at E9.5 (*SI Appendix*, Fig. S7*A*), and are both down-regulated in the otic vesicle epithelial cells and otic geniculate ganglion cells at E11.5 (*SI Appendix*, Fig. S7*B*), and are close to undetectable in the inner ear sensory and non-sensory epithelial cells at E13.5 (*SI Appendix*, Fig. S7*C*). At P2, *Trim71* and *Sall4* are near-undetectable in cochlear HCs, SCs (PC/DC), and surrounding non-epithelial cells (*SI Appendix*, Fig. S7*D*). However, *Sall2* and *Sall3* exhibited robust expression in pillar and Deiter’s cells at P2 (SI Appendix, Fig. S7*D*), suggesting a potential role for them as SC-specific genes. This notion is further corroborated by the observed downregulation of both genes in TRIM71-overexpressing organoids relative to controls (SI Appendix, Fig. S6).

To gain insight into how TRIM71 reprograms SCs (and GER cells) into proliferative, progenitor-like cells, we conducted gene ontology (GO) enrichment analysis of genes differentially expressed between TRIM71-overexpressing and control cells. We found that terms such as stemness, cell proliferation and epithelial-mesenchymal transition were associated with genes that were upregulated in TRIM71-overexpressing cells, whereas terms involved in cell differentiation, such as TGFβ signaling and extracellular matrix were downregulated for clusters 0 (*SI Appendix*, Fig. S8*A*), 1, 3, 5, 8 (*SI Appendix*, Fig. S8 *B* and *C*), 2, 4, 6, 7 (*SI Appendix*, Fig. S8*D*).

To investigate the mechanism by which FST coactivation with TRIM71 promotes HC formation in cochlear organoid culture, we performed a GO enrichment analysis comparing TRIM71-overexpressing and FST+TRIM71-overexpressing cell clusters. We found that genes involved in pathways and biological processes associated with TGF-β signaling and epithelial-to-mesenchymal transition were significantly enriched among FST+TRIM71-downregulated genes (Fig. 4*E*). Consistent with these observations, RT-qPCR experiments showed that mRNA expression levels of the mesenchymal marker genes *Twist1*, *Scx*, and *Mmp14* were significantly increased in TRIM71-overexpressing organoids relative to control organoids and that this up-regulation was attenuated in organoids that co-expressed FST+TRIM71 (Fig. 4*F*).

By contrast, pathways and biological processes associated with HC formation such as WNT signaling pathway, sensory organ morphogenesis, inner ear development, and inner ear receptor cell differentiation were significantly enriched among FST+TRIM71 up-regulated genes (Fig. 4*E*). Collectively, these data suggest that FST may be critical to limit aberrant activation of mesenchymal genes in TRIM71-overexpressing SCs and GER cells, which may enhance the rate of HC formation by preventing the conversion of SCs/prosensory cells into mesenchymal cells.

Previous studies have found that regulatory sequences of HC genes become less accessible as cochlear SCs begin to mature (28). To determine whether TRIM71 and/or FST+TRIM71-overexpressing increase chromatin accessibility at HC-specific gene loci, we conducted ATAC-seq (Assay for Transposase Accessible Chromatin sequencing) experiments with control, FST, TRIM71, and FST+TRIM71-overexpressing cochlear organoids. We found that ATAC-seq peaks around transcription start sites (TSS) were increased in FST+TRIM71-overexpressing samples compared with control, FST or TRIM71-overexpressing samples (Fig. 5, *A* and *B*), indicating genome-wide higher chromatin accessibility. We next examined transcription factor motif enrichment in differentially accessible chromatin regions between control, FST, TRIM71, and FST+TRIM71. Motifs for GATA transcription factors (GATA1, GATA2, GATA4, GATA6) and ZIC3 were significantly enriched for both TRIM71 and FST+TRIM71-overexpressing samples, and GATA3 was only enriched in the FST+TRIM71-overexpressing sample (Fig. 5, *C* and *D*). GATA and ZIC transcription factors are key regulators of human primed pluripotency (39, 40). GATA3 is required for cochlear prosensory development and the functional maturation of inner HCs (41, 42).

**Fig. 5.**
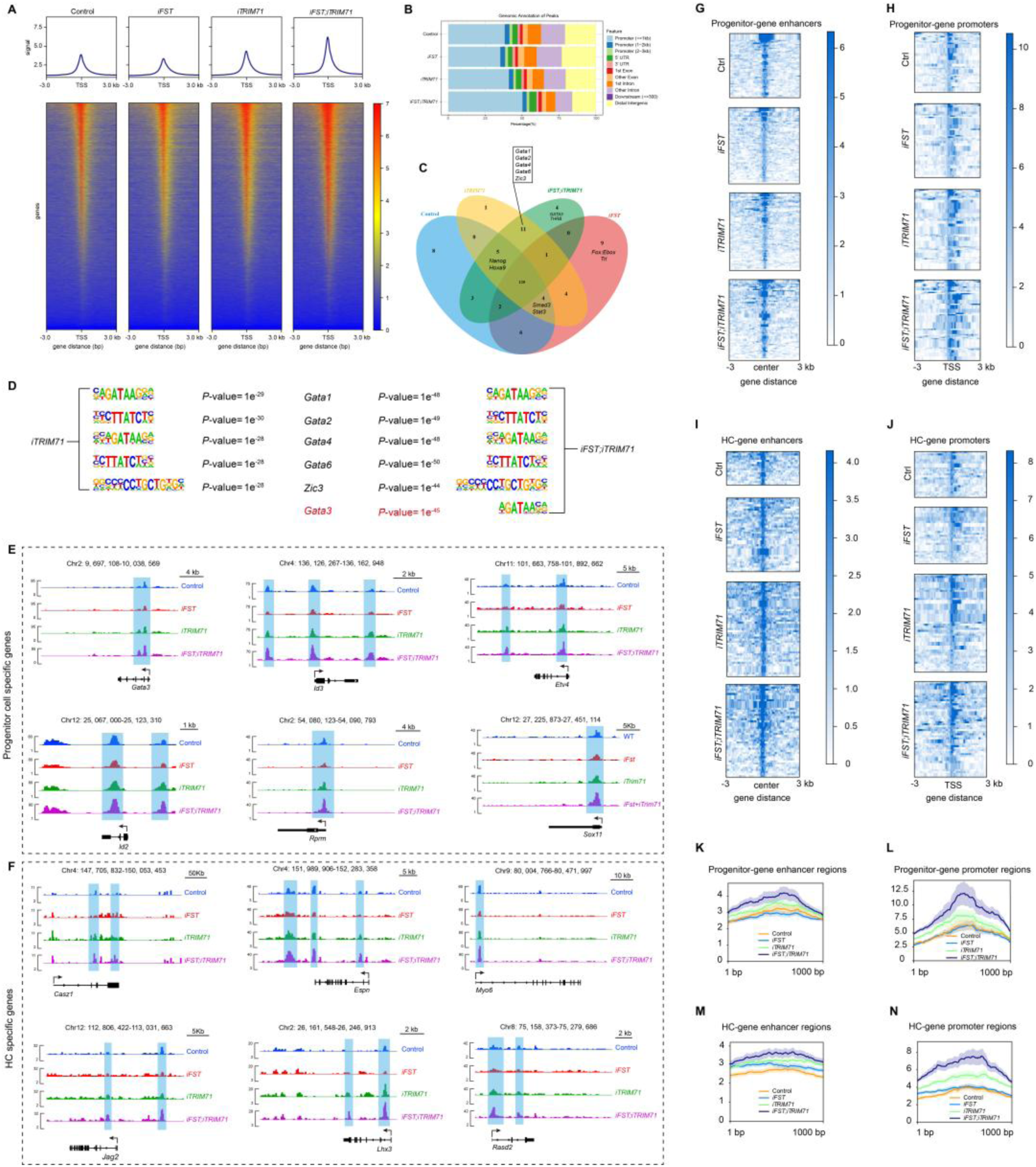
Combined FST and TRIM71 overexpression enhances the chromatin accessibility at HC/progenitor gene promoters and enhancers in cochlear organoid culture. Cochlear epithelial cells from P5 control, *iFST*, *iTRIM71*, and *iFST;iTRIM71* tg mice were used to establish organoid cultures. Cultures were analyzed after 10 days of expansion. To induce expression of FST, TRIM71, or both, Dox was added to the medium at the start of culture. (*A*) The ATAC-seq signal enrichment around transcription start sites in control, FST, TRIM71, and FST+TRIM71 cochlear organoids overexpressing. Peaks representing open chromatin are shown for a ∼6 kb region of the mouse genome near the gene locus. (*B*) Genomic annotation of peaks. (*C*) Venn diagram depicting the overlap among the sets of transcription factor binding motifs that were enriched in the differentially accessible regions from control, FST, TRIM71, and FST+TRIM71-overexpressing cochlear organoids. (*D*) Inspection of the overlapping motif sets reveals that *Gata1*, *Gata2*, *Gata4*, *Gata6* and *Zic3* motifs are commonly enriched in the intersection shared by the TRIM71 and FST+TRIM71 conditions, whereas the Gata3 motif is specifically enriched in the FST+TRIM71-unique set. (*E-F*) ATAC-seq peaks for progenitor-specific genes (*Gata3*, *Id2*, *Id3*, *Rprm*, *Etv4*, *Sox11*) and HC-specific-genes (*Casz1*, *Jag2*, *Espn*, *Lhx3*, *Myo6*, *Rasd2*). (*G-J*) Chromatin structure (ATAC) was compared at progenitor/HC gene promoters and enhancers in control, FST, TRIM71 and FST+TRIM71-overexpressing cochlear organoids. (*K-N*) Comparison of average signal profiles (1000 bp) for each group indicate that progenitor/HC gene promoters and enhancers gain accessibility after overexpression of TRIM71, with is highest in FST+TRIM71-overexpressing samples.

To determine whether regulatory sequences of progenitor and/or HC genes are more accessible in TRIM71-overexpressing samples, we analyzed ATAC-seq peak frequency at recently identified progenitor- and HC-specific promoters and enhancers (34). Chromatin accessibility at these gene loci was higher in TRIM71 overexpressing organoids compared with control, which reaches highest in FST+TRIM71-overexpressing samples, as illustrated by the genome browser tracks for progenitor-cell specific genes (*Gata3*, *Id2*, *Id3*, *Rprm*, *Etv4*, *Sox11*) and HC-specific genes (*Casz1*, *Jag2*, *Espn*, *Lhx3*, *Myo6*, *Rasd2*) (Fig. 5, *E* and *F*). Heatmaps also revealed that in FST+TRIM71-overexpressing samples, chromatin accessibility at promoters and enhancers of progenitor (Fig. 5, *G* and *H*) and HC genes (Fig. 5, *I* and *J*) was higher than the other samples. Consistent with the above data, the average signal profiles (1000 bp) for each group further validated the positive effect of FST and TRIM71 on chromatin accessibility of progenitor cell and HC specific genes (Fig.5, K-N). In summary, these data suggest that FST+TRIM71-overexpression promotes expression of progenitor and HC-specific genes, at least in part, by increasing chromatin accessibility at their promoters and enhancers.

### The RNA Binding Domain of TRIM71 is Essential for *Sall4* Activation in Cochlear Organoid Culture

TRIM71 has a dual function as an RNA-binding protein and a ubiquitin ligase (43, 44). We recently found that TRIM71’s ubiquitin ligase activity is dispensable for SC plasticity; however, its RNA-binding activity is essential (15). To elucidate whether the RNA-binding activity is required for *Sall4* activation during HC formation, we overexpressed wild-type TRIM71 or TRIM71 lacking the RNA-binding domain (ΔNHL) in cochlear organoid culture (SI Appendix, Fig. S9A, and *B*), and differentiated the organoids in differentiation media containing CHIR99021 together with the γ-secretase and Notch inhibitor LY411575 as reported previously(24, 25). We found fewer Atoh1-nGFP⁺ cells in ΔNHL-overexpressing organoids than in TRIM71-overexpressing organoids (SI Appendix, Fig. S9*C*). Transcriptional analysis also revealed that the expression of early HC-genes (*Atoh1*, *Pou4f3*, *Gfi1*, and *Myo7a*) and progenitor genes (*Hmga2* and *Sall4*) was significantly down-regulated in ΔNHL-overexpressing organoids compared with TRIM71-overexpressing control (SI Appendix, Fig. S9*D*). This suggests that the RNA-binding domain of TRIM71 is required for inducing Sall4 expression in cochlear organoid culture.

### *Sall4* is Required for HC Regeneration in Cochlear Organoid and Explant Culture in Response to TRIM71 Overexpression

To investigate whether *Sall4* is required for HC formation, we knocked down *Sall4* in organoid cultures established from *Atoh1-nGFP* tg mice using lentiviral transduction (Fig. 6*A*). A blue fluorescent protein (BFP) reporter was used to monitor lentiviral infection in organoids. Among the three short hairpin RNA constructs tested, *shRNA-Sall4-2#* exhibited the highest knockdown efficiency for *Sall4* and was therefore selected for subsequent experiments (Fig. 6*B*). Loss of *Sall4* significantly reduced the number of Atoh1-nGFP⁺ organoids (Fig. 6*C*) and concurrently downregulated the mRNA levels of *Atoh1*, *Pou4f3* and *Gfi1* relative to control (Fig. 6*D*).

**Fig.6.**
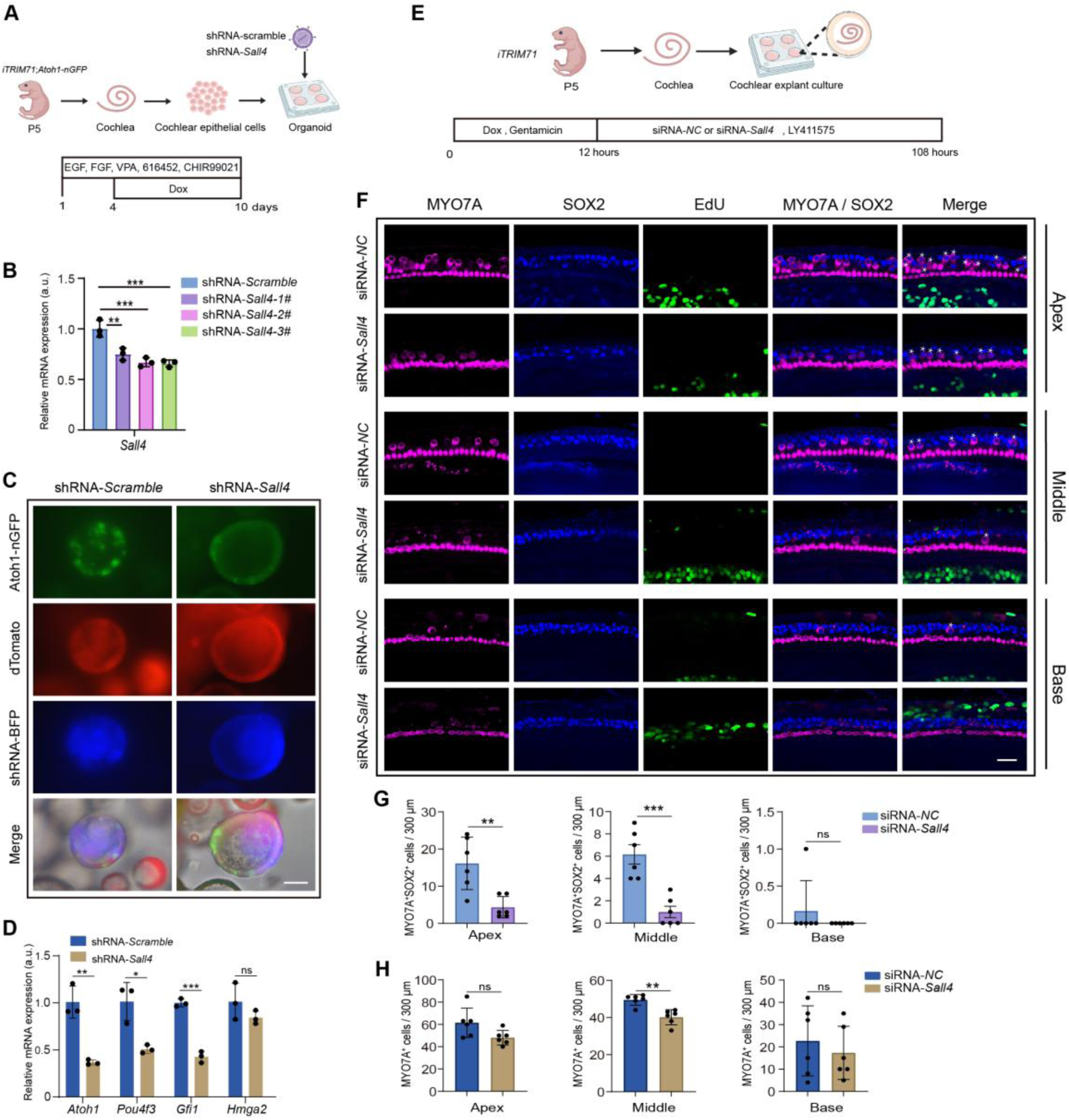
Sall4 is necessary for HC formation in cochlear organoid and explant culture. (*A*) Experimental scheme for (*B-D*) Created in BioRender. (*B*) RT-qPCR analyzing the knockdown efficiency of *Sall4* after infected shRNA-scramble, shRNA-*Sall4*-1#, shRNA-*Sall4*-2#, shRNA-*Sall4*-3#. (*C*) Low-power bright-field, green (Atoh1-nGFP), red (tdTomato), and blue fluorescent (BFP) images of organoid cultures in shRNA-Scramble and shRNA-*Sall4*. Scale bars=200 μm. (*D*) RT-qPCR analyzing HC-specific mRNA expression (*Atoh1*, *Pou4f3*, *Gfi1*) and Hmga2 in control (shRNA-scramble) and *Sall4* knockdown (*shRNA*-*Sall4*) organoids after 10 d of expansion (n=3 animals, two independent experiments). (*E*) Experimental scheme for (*F-H*). P5 cochlear explants from *iTRIM71* mice were dissected and cultured on SPI-Pore membrane filters, and gentamicin was used to damage HCs. After 12 h of culture, gentamicin was removed. Then siRNA-*NC*, siRNA-*Sall4,* including LY411575, were added to the culture medium for an additional 4 days. Created in BioRender. (*F*) Confocal images of cochlear sensory epithelia after 4 days of culture stained for MYO7A (magenta), SOX2 (blue), and EdU (green). White asterisks label newly formed MYO7A^+^SOX2^+^ HCs. Scale bars=40 μm. (*G-H*) Quantification of newly formed HCs (*G*) and total number of HCs (*H*) in control and *Sall4-* knockdown cochlear explants at the apex, middle, and base regions after TRIM71 overexpression (n=6 animals, three independent experiments). Two-tailed, unpaired Student’s *t* test was used to calculate *P* values for *D*, *G*, *H*. *\*P<0.05*, *\*\*P<0.01*, *\*\*\*P<0.001*. n.s., not significant.

Next, we evaluated whether the function of *Sall4* is necessary for TRIM71-mediated HC regeneration in cochlear explants. To do so we knocked down *Sall4* expression using small interfering RNAs (siRNAs) in P5 TRIM71-overexpressing cochlear explants. As control, P5 TRIM71 overexpressing explants received non-specific siRNA. To induce HCs regeneration, explants were exposed to gentamicin overnight and treated with Notch inhibitor (Fig. 6*E*). We found that *Sall4* knockdown significantly reduced the number of regenerated HCs (MYO7A^+^SOX2^+^) in the apical and middle portion of TRIM71 overexpressing cochlear explants (Fig. 6 *F* and *G*). Consistent with such decline in regenerative capacity, *Sall4* knockdown also reduced the total number of HCs (MYO7A^+^) (Fig. 6 *F* and *H*), in the middle portion of TRIM71 overexpressing explants. There were no EdU^+^MYO7A^+^SOX2^+^ cells in response to TRIM71 overexpression (Fig. 6*F*), suggesting that newly formed HCs in TRIM71-overexpressing cochlear explants originated from non-mitotic cells.

### SALL4 Overexpression Promotes Cochlear Organoid Proliferation but Inhibits HC Formation

SALL4 functions as a reprogramming factor for iPSCs and has been shown to promote tissue regeneration (37). Our previous study showed that *Trim71* deficiency impairs HC formation in organoid cultures established from E13.5 cochlear epithelial cells (15). To investigate whether SALL4 is sufficient to rescue the lack of HC formation in *Trim71*-deficient organoids, we overexpressed HA-tagged SALL4 in organoids established with cochlear epithelial cells obtained from P5 wildtype and *Trim71^Δ/Δ^* mice (SI Appendix, Fig. S10*A*). We overexpressed HA-tagged SALL4 using a lentivirus that also co-expressed mCherry, which enabled us to track infected cells. As for control, wild type and *Trim71*-deficient organoids were infected with lentivirus expressing HA-tag only (SI Appendix, Fig. S10*B*). Immunostaining with HA antibody showed, as expected, diffuse HA peptide expression throughout the cell in organoids infected with control lentivirus and nuclear HA expression in organoid cultures infected with HA-tagged SALL4 expressing lentivirus (SI Appendix, Fig. S10*C*). We found that at 10 days of expansion, the organoid diameter was significantly larger in SALL4-overexpressing control and *Trim71^Δ/Δ^* organoids compared to organoids that overexpressed HA-tag only (SI Appendix, Fig. S10, *D and E*), and the number of organoids increased by about 1.5-fold (SI Appendix, Fig. S10*F*). After 4 days of differentiation, we found that the mRNA levels of HC-specific genes *Atoh1* and *Myo7a* were downregulated in *Trim71^Δ/Δ^* mice. In addition, we found that SC-gene *S100a1* and progenitor-gene *Hmga2* were lower expressed in Trim71-deficient organoids than control organoids. Furthermore, we found that SALL4 overexpression further reduced the expression levels of *Atoh1*, *Myo7a*, and *S100a1* in both control and *Trim71^Δ/Δ^* organoids compared with HA overexpression (SI Appendix, Fig. S10, *G* to *J*). Based on these results, we propose that SALL4 primes SCs to acquire a progenitor-like state; however, SALL4 overexpression prevents further differentiation of progenitor-like cells into HCs in cochlear organoid culture.

## Discussion

We recently discovered that TRIM71, an RNA-binding protein and ubiquitin ligase, is a potent activator of cochlear SC plasticity, enhancing SC division and HC formation in vitro (15). However, many questions remained, including the transcriptional and epigenetic mechanisms through which TRIM71 promotes SC plasticity and HC formation, as well as whether TRIM71 promotes HC regeneration *in vivo*.

By utilizing a Dox-inducible human TRIM71 transgenic mouse line (*iTRIM71*), we show that TRIM71 activation enhances HC formation in vitro and in vivo. However, it is important to note that reactivation of TRIM71 alone is insufficient to stimulate SC proliferation or to induce HC formation. Rather, TRIM71 greatly enhances the “responsiveness” of SCs to mitogens and HC-fate-inducing factors by reprogramming SCs into HC progenitor-like cells. This is also illustrated in our in vivo experiments, where we find that TRIM71 overexpression in the damaged cochlea enhances spontaneous HC regeneration but does not in the undamaged cochlea. Interestingly, following HC injury, *Trim71* is rapidly upregulated in cochlear epithelial cells. Future studies will determine whether *Trim71* is part of the machinery that triggers spontaneous HC regeneration in the immature cochlea.

In addition to requiring HC-fate-inducing signals such as HC damage or Notch inhibition, inhibition of TGFβ-type signaling, achieved through FST overexpression, is critical for TRIM71-mediated enhancement of HC formation in P5 cochlear organoid and explant culture. FST overexpression has alone does not affect HC formation, indicating its role as a cofactor. How does FST facilitate HC formation in response to TRIM71 overexpression? Our scRNA-seq data indicate that TRIM71 is a potent driver of the reprogramming of cochlear SCs into progenitor-like cells. However, in the absence of FST, TRIM71 fails to induce early HC genes but instead activates genes associated with epithelial-to-mesenchymal transition, which may, if prolonged, convert SCs into mesenchymal-like cells. We propose that FST overexpression promotes HC regeneration by counteracting the aberrant activation of the epithelial-to-mesenchymal transition induced by TRIM71 overexpression.

In addition, our ATAC-seq experiment suggests that co-activation of FST and TRIM71 increases accessibility of HC- and progenitor-specific promoters and enhancers and selectively increases chromatin accessibility at GATA3-binding sites. The transcription factor GATA3 is a key regulator of early otic and cochlear neurosensory development and is essential for inner HC maturation (42). Co-activation of GATA3 with ATOH1 facilitates the conversion of adult cochlear SCs into HCs (45). How the combined activation of FST and TRIM71 increases chromatin accessibility at HC and progenitor gene loci, and how the functions of TRIM71 and GATA3 may intersect in HC regeneration, remain to be determined. The *Gata3* motif was only significantly enriched in FST+TRIM71-overexpressing organoids, suggesting that GATA3 may be a key effector of FST and TRIM71 during HC regeneration.

TRIM71 is a potent post-transcriptional repressor and has been shown to promote the reprogramming of human induced pluripotent stem cells by antagonizing pro-differentiation signals and factors (46). However, we find that in addition to such repressive function, TRIM71 is a potent activator of early otic progenitor genes, including Sall4. SALL4 belongs to the core pluripotency network that represses somatic cell differentiation and promotes reprogramming of somatic cells into pluripotent stem cells (47, 48). *Sall4*, a member of the evolutionarily conserved spalt-like gene family (*Sall1-4*), is essential for early embryonic development and stemness (32). In the chick inner ear, Sall4 regulates otic placode gene expression and is essential for placode invagination (49).

Our study revealed that *Sall4* shows a similar expression pattern to *Trim71*. *Sall4* is co-expressed with *Trim71* in early otic progenitors and later in cochlear epithelial cells. Using cochlear organoid and explant cultures, we find that shRNA and siRNA-mediated knockdown of *Sall4* attenuates HC formation and HC regeneration. However, while SALL4 overexpression promotes SC proliferation in cochlear organoid culture, it interferes with HC formation, suggesting that SALL4 is involved in reprogramming SCs into progenitor-like cells but inhibits their further differentiation into HCs.

Here, we provide evidence that the decline in murine cochlear SC plasticity, at least in part, correlates with the downregulation of *Trim71-Sall4*. We establish a regulatory link between cochlear *Trim71* and *Sall4* expression levels *in vitro* and *in vivo*. Our data indicate that *Sall4* is a downstream target of TRIM71, which is essential for HC regeneration. It will be of interest to further explore whether overexpression of *Sall4* and *Trim71,* together with *Atoh1*, can synergistically promote HC regeneration at P7 or even in the adult stage.

## Acknowledgments

We thank the members of the Li Laboratory for their help and advice throughout this study. We thank Mr. Guo at the FIST Sharing Platform of Xi’an Jiaotong University for their assistance with confocal imaging analysis.

## Fundings

National Natural Science Foundation of China grant 32570978 (X.J.L.)

Natural Science Basic Research Plan in Shaanxi Province of China, No.2024JC-ZDXM-43 (X.J.L.)

Fundamental Research funds for the Central Universities (X.J.L.)

National Institute on Deafness and Other Communication Disorders, grant R01DC011571 (A.D.)

David M. Rubenstein Fund for Hearing Research (A.D.)

## Author contributions

Conceptualization: X.J.L., A.D., L.L., H.H.

Data curation: X.Y.L., L.L., X.Y.L., Y.F.W.

Methodology: X.J.L., L.L., S.X.M., R.F.Q., Y.Y.X., Q.H., H.M., B.S., X.Y.M., J.L., S.H.D.

Visualization: X.J.L., L.L., H.H., X.Y.L., Y.H.W., Y.Z.,

Investigation: X.J.L., L.L.,

Supervision: X.J.L., A.D., W.K.

Writing—original draft: X.J.L., A.D., L.L., H.H.,

Writing—review & editing: X.J.L., A.D., L.L., H.H.,

