## Supplemental Data 1 for "The RNA-binding protein TRIM71 Promotes Supporting Cell Reprogramming and Hair Cell Regeneration by Activating *Sall4*"

**Affiliations**

*These authors contributed equally

**This PDF file includes:**

Supporting text

Figures S1 to S10

Tables S1 to S4

**Other supplementary materials for this manuscript include the following:**

Datasets S1

### Supplementary Materials and Methods

#### Mouse breeding and genotyping

All experiments and procedures were approved by Xi'an Jiaotong University Animal Care and Use Committee protocol (XJTUAE2025-199). *TRE-FST-288* transgenic mice were obtained from S.-J. Lee (the Jackson Laboratory, Farmington) (1). *R26rtTA*M2* (no. 006965) mice were purchased from the Jackson Laboratory (Bar Harbor, ME). The *Atoh1-nGFP* tg mice were obtained from J. Johnson (University of Texas Southwestern Medical Center, Dallas) (2). *Trim71* floxed mice (RRID: MGI:7661049) were provided by Waldemar Kolanus, Universität Bonn. $Pou4f3^{DTR/+}$ (no. 028673) mice were purchased from the Jackson Laboratory (Bar Harbor, ME). All the transgenic mouse lines were shared by Angelika Doetzlhofer's lab at Johns Hopkins University. *iTRIM71* mice were obtained from Cyagen Company. Both male and female mice were used in this study. The genotyping primers are listed in Table 1.

#### Organoid culture

Inner ears were dissected in cold HBSS (Corning, no. 21-023-CV), and cochlear epithelial cells were isolated by incubating the cochlear tissues with collagenase (1 mg/mL; Worthington, no. LS004214) and dispase (1 mg/mL; ThermoFisher, no. 17105041) for 8 min at room temperature. DMEM/F12 (Corning, no. 10-092-CV) supplemented with 10% serum was used to terminate enzymatic digestion.. Finally, TrypLE solution (ThermoFisher, no. 2604013) was used to further dissociate the cochlear epithelium into single cells. The cell pellets were resuspended in expansion medium and then mixed with an equal volume of Matrigel (Corning, no. 356231), and plated into the middle of prewarmed 4-well plates. The culture medium was changed every other day. The components of the expansion and differentiation medium were as described previously (3-5).

#### Quantification of organoid number, organoid diameter, and GFP reporter expression

Low-power bright-field and fluorescence micrographs of organoid cultures were acquired using an IBE2000 microscope equipped with 4× and 10× objectives. For organoid size assessment, the organoid diameter was measured in three to four randomly selected fields per culture using ImageJ software, and the mean diameter was recorded as a single data point. For each genotype and/or treatment condition, a minimum of three biologically independent organoid cultures were established and analyzed. At least two independent experimental replicates were performed and evaluated. All measurements and data analyses were performed with the investigators blinded to genotype and/or treatment assignment.

#### Cochlear explant culture

Cochlear sensory epithelia including innervating neurons (cochlear explants) from stage P2 or stage P5 mice were isolated by microdissection, and cultured on SPI-Pore membrane filters (Structure Probe, no. E1013-MB) in N-2 supplement (1×; Gibco, no.17502-048), B-27 supplement (1×; Gibco, no.12587-010), EGF (5 ng/ml; Sigma-Aldrich, no.SRP3196), Ampicillin (10 mg/ml; Beyotime, no. ST008), LY411575 (5 mM; Sigma-Aldrich, no. SML050), HEPES (10 mM; Beyotime, no. C0215) and Glutamax (1×; Gibco, no.35050-061). Gentamicin sulfate salt (25 μg/ml; Beyotime, no. ST2398) was used to ablate HCs. Doxycycline (0.5 μg/ml; Sigma-Aldrich, no. D9891) was used to induce FST and/or TRIM71 overexpression. To trace dividing cells, EdU (3 μM; MCE, HY-118411) was added to the medium. In all experiments, the culture media were changed every other day, and cultures were maintained for a total of four days.

**RNA extraction and RT-qPCR**

Cochlear organoids were recovered from Matrigel using Cell Recovery Solution (Corning, no. 354253), then washed three times with 1×PBS. Cochlear epithelia were isolated from cultured cochlear explants or freshly harvested inner ear tissue using dispase/collagenase treatment. Total RNA was extracted with TRIZOL reagent (Takara, no. 9109) and reverse-transcribed into cDNA using a PrimeScript RT reagent kit (Vazyme, no. R223). RT-qPCR was performed with SYBR Master Mix(Vazyme, no. Q312) and gene-specific primer sets on a Real-Time PCR Detection System (Agilent Technologies, Mx3000P). Each PCR was performed in triplicate. The ribosomal gene *Rpl19* was used as an endogenous reference gene. The RT-qPCR primers were listed in Table 2.

**Single-cell RNA sequencing**

Organoids were established with the cochlear epithelial cells of P5 mice. After 10 days of expansion culture, the organoids were harvested and enzymatically dissociated into a single-cell suspension. Cell viability and concentration were determined using Countstar Rigel (Alit Biotech). To enrich for viable cells, dead cells were subsequently removed using the Dead Cell Removal Kit (Miltenyi Biotec, no. 130-090-101). Finally, cell pellets were resuspended in PBS containing 0.04% BSA to achieve a final target concentration of 700-1200 cells/μl. The single-cell suspension was loaded onto a 10x Genomics Chromium Chip (v4, PN: 1000690) and barcoded using a 10x Genomics Chromium Controller. Following cell barcoding, RNA was reverse-transcribed, amplified, and processed into sequencing libraries using the 10x Library Construction Kit (v4, PN: 1000694), with all procedures performed in accordance with the manufacturer's recommended protocols. Sequencing was performed on an Illumina NovaSeq using 150-bp paired-end reads at Novogene Bioinformatics Technology Co., Ltd. (Tianjin, China). Raw data were processed with Cell Ranger (9.0.1) to demultiplex reads and generate gene-by-cell UMI count matrices. Reads were aligned to the mouse GRCm39-2024-A reference genome.

**Single-cell RNA data analysis**

Cell Ranger output data was performed with the Seurat (v5.1.0) pipeline (6). First, cells were filtered based on the number of detected genes (1,000–7,000 genes per cell). Cells with more than 10% mitochondrial gene expression were excluded. After the pre-processing step, data were log-normalized, and the top 2,000 highly variable genes were identified with the vst method with default settings. We scaled the datasets to avoid the dominance of highly expressed genes and used the data as input for PCA to reduce dimensionality. The Harmony-corrected (v1.2.3) (7) embeddings were then used for downstream analysis. The shared nearest neighbor graph was constructed using 40 nearest neighbors based on the first 25 Harmony-corrected dimensions. The shared nearest-neighbor modularity optimization-based clustering algorithm was used to identify clusters with resolution 0.4. UMAP was used to visualize the scRNA-seq clustering results and metadata information. To determine cell identities for each cluster, we used the FindAllMarkers function in the Seurat package with the following parameters: only.pos=TRUE, min.pct=0.25, logfc.threshold=0.25. Adjusted Bonferroni-corrected P-values of 0.05 were used for multiple testing correction. We determined cell identities by comparing cluster-specific DEGs with published canonical marker genes. Differentially expressed gene (DEG) analysis was performed using the FindAllMarkers function with parameters: min.pct = 0.1, logfc.threshold = 0.25. Genes with $|log2FC| > 1$ and adjusted p-value $< 0.05$ were considered significantly differentially expressed. Gene Ontology (GO) enrichment and Kyoto Encyclopedia of Genes and Genomes (KEGG) analysis were performed using clusterProfiler (v4.14.4) (8) with significantly differentially expressed genes.

**ATAC-seq library preparation and sequencing**

To prepare single-cell suspensions, cochlear organoids were recovered from Matrigel using cell recovery solution (Corning, 354253). Subsequently, the collected organoids were digested with TrypLE Express (ThermoFisher, no. 2604013) at 37°C for 5 min, followed by neutralization with medium containing 10% FBS. The mixture was gently and repeatedly pipetted to mechanically dissociate the organoids into a single-cell suspension, which was then counted. A total of 100,000 cells were utilized as input and subjected to ATAC-seq library construction using the ATAC-Seq Library Prep Kit (Vazyme, no. TD711) according to the manufacturer’s instructions. Finally, all ATAC-seq libraries were sequenced on the NovaSeq X Plus Series (PE150).

**ATAC-seq analysis**

Adapter contamination and low-quality bases were removed from the raw sequencing reads using Trim Galore (v. 0.6.6), and the cleaned reads were subjected to quality control with FastQC (v. 0.11.9). The high-quality reads were aligned to the mouse reference genome

(GRCm38/mm10) using Bowtie2 (v. 2.5.4) with the --very-sensitive and --no-unal parameters. Peak calling was performed using the MACS2 algorithm (v. 2.2.7.1) with a q-value threshold of 0.05. The resulting peaks were annotated with genomic features (e.g., promoters, exons, introns, and intergenic regions) using ChIPseeker (v. 1.40.0), which was also employed to generate stacked bar charts summarizing the annotation composition. Motif enrichment analysis was performed with HOMER (v. 1.80.2) to identify overrepresented DNA sequence motifs within the peak sets. Coverage tracks were visualized with Integrative Genomics Viewer (IGV) (v. 2.11.9); TSS enrichment heatmaps and average signal profiles were generated with deepTools (v. 3.5.2). A Venn diagram was constructed with VennDiagram (v. 1.7.3) to illustrate overlaps among different peak sets.

To investigate changes in chromatin accessibility of HC- and progenitor cell-specific genes across control, FST, TRIM71, and FST+TRIM71-overexpressing cochlear organoids, we analyzed the enhancer and promoter peak regions for these genes (Datasets S1). We calculated the signal intensity in the specific regions using 'computeMatrix', and converted the data into heatmaps using 'plotHeatmap'. For enhancer regions, we selected signals within 3 kb upstream and downstream of the peak center as the reference point. For promoter regions, we selected signals within 3 kb upstream and downstream of the transcription start site (TSS).

**Immunohistochemistry**

Cochlear organoids and explants were fixed with 4% (vol/vol) paraformaldehyde for 30 min, followed by three washes in PBS. Then cochlear organoids and explants were incubated with 0.5% (vol/vol) TritonX-100/ 10% (vol/vol) FBS in 1×PBS for 30 min to permeabilize cells. The primary antibodies were diluted in 1×PBS and incubated overnight at 4 °C, and the secondary antibodies were also diluted with 1×PBS and incubated at room temperature for 2 hours in the dark. The antibodies used were listed in Table 3.

**Cell proliferation**

EdU was added to culture media at a final concentration of 3 μM. Cell proliferation was detected by quantifying the percentage of EdU-incorporated cells using Click-iT Edu Alexa Fluor 488 imaging Kit (Vazyme, no. A411) and Click-iT Edu Alexa Fluor 555 imaging Kit (ThermoFisher, no. C10338) following the manufacturer's specifications.

**Lentiviral vectors and production**

To overexpress SALL4, we constructed a lentiviral overexpression vector that, under the control of the CMV promoter, expresses the human SALL4 coding sequence (CDS) that is fused to a 3×HA tag and T2A-linked mCherry fluorescent protein, which we used to monitor the infection of lentivirus and expression of SALL4. The plasmids encoding wild-type TRIM71 and ΔNHL-TRIM71 were derived from plasmids used in our previous work (3). All plasmids used for *Sall4* knockdown were constructed from the PLKO.1-TagBFP-Puro vector (FENGHUISHEWU, no. ZT1579). The PLKO.1-TagBFP-Puro vector was linearized using AgeI-HF (New England Biolabs, no. R3552S) and EcoRI-HF (New England Biolabs, no. R3101S) restriction enzymes. Primers were annealed to form an insert fragment with AgeI-HF and EcoRI-HF sticky ends at both ends, which was then ligated to the linearized vector using T4 ligase. The sequence of shRNAs are as follows: shRNA-Scramble-BFP, 5′-cctaaggttaagtcgccctcgctcgagcgagggcgacttaaccttagg-3′; shRNA-Sall4-1#-BFP, 5′-gaagcctttcgtgtgtaacatctcgagatgttacacacgaaaggcttc-3′; shRNA-Sall4-2#-BFP, 5′-cagcccacctttgtcaaagttctcgagaactttgacaaaggtgggctg-3′; shRNA-Sall4-3#-BFP, 5′-gcccacctttgtcaaagttgactcgagtcaactttgacaaaggtgggc-3′. For packaging, 293T cells (ATCC, no. CRL-3216, RRID: CVCL_0063) were transfected with 10 µg of transfer plasmid, 2 µg of pMDLg/pRRE (Addgene, no. 12251), 2µg of pRSV-Rev (Addgene, no.12253), and 2 µg of pCMV-VSV-G (Addgene, no.8454). After 48-72 h of culture, the supernatant was collected, centrifuged at 500 g for 10 min at 4°C, and filtered through a 0.45µm PES membrane filter to remove cell debris. Lentiviral particles were concentrated using ultracentrifugation at 90,000 g for 1.5 hours at 4°C. The pellet was resuspended in 100 µl of DMEM/F12 medium overnight at 4°C and stored at −80°C.

**siRNA Transfection**

Cochlear explants were transfected with siRNAs targeting Sall4 (synthesized by Tsingke Biotechnology) using AccuFect RNAi Transfection Reagent (ACCURATE BIOTECHNOLOGY, AG51017) for 24 hours, following the manufacturer's instructions. The sequences of siRNAs are listed in Table 4.

**Statistical analysis**

All experimental findings were validated through at least two independent replicate experiments. The sample size (n) denotes the number of animals analyzed per group, with each animal serving as an independent biological replicate. Animals were assigned to control or experimental cohorts based on their genotype and/or treatment regimen. To minimize potential bias, both blinding and randomization were implemented during data acquisition and analysis. Statistical analyses were performed using GraphPad Prism 10 software. Detailed specifications for each experiment—including sample sizes, statistical tests applied, and corresponding P values—are provided in the respective figure legends. Statistical significance was defined as $*P \leqslant 0.05$, $**P<0.01$, $***P<0.001$, and $****P<0.0001$. All error bars indicate standard deviation (SD) calculated from biological replicates.

### Figures S1 to S10

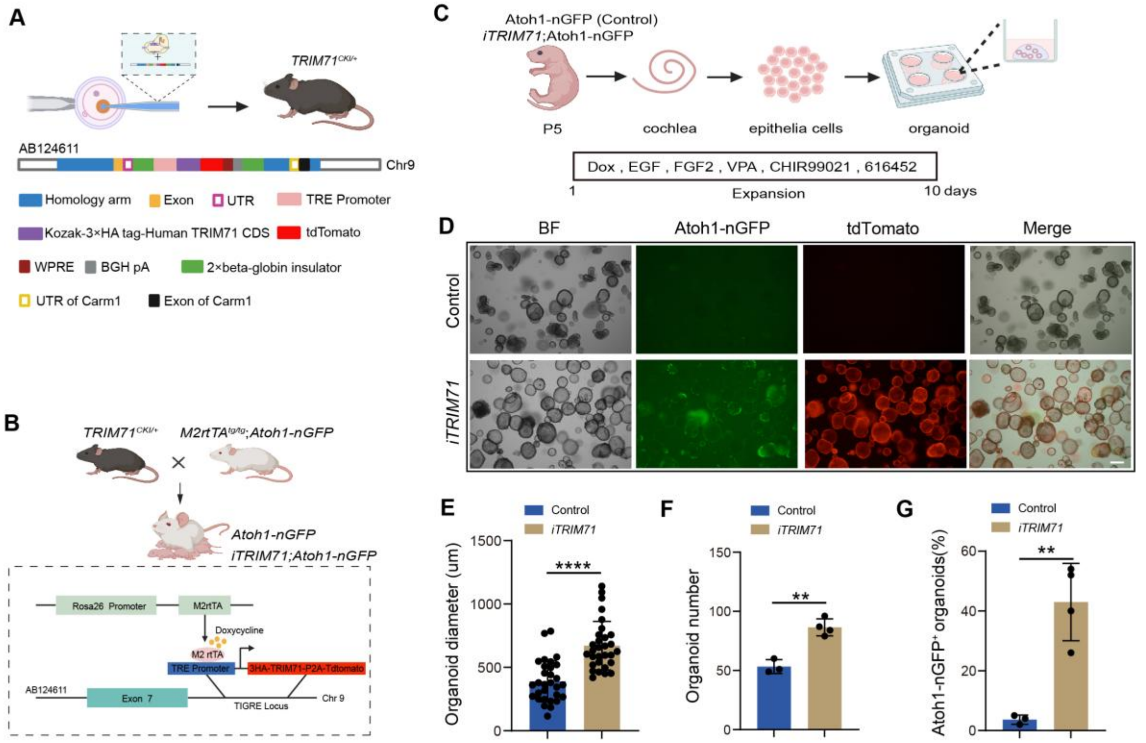


**Fig. S1. Generation and validation of the TRIM71 transgenic mouse model.** (*A*-*B*) Schematic of the *iTRIM71* transgenic mouse model. A transgene expressing 3×HA-tagged human TRIM71 CDS under the control of the TRE promoter was combined with an M2 reverse tetracycline transactivator (M2rtTA) transgene driven by the ubiquitously expressed Rosa26 promoter to enable global TRIM71 expression. Created in BioRender. (*C*) Experimental scheme for (*D* to *G)*. Created in BioRender. (*D*) Low-power bright-field, green (Atoh1-nGFP) and red (tdTomato) images of organoid cultures. Atoh1-nGFP marks nascent HCs. Scale bar, 200 µm. (*E*) Organoid diameters in (D) (organoid number = 30, 30 organoid from three independent experiments were quantified). (*F*) Organoid number in (*D*) (n=3 in WT group, n=4 in *iTRIM71* group). (*G*) Percentage of Atoh1-nGFP+ organoids in (D) (n=3 in WT group, n=4 in *iTRIM71* group). Two-tailed, unpaired Student's *t* test was used for *E-G*. $****P < 0.0001$, $**P < 0.01$.

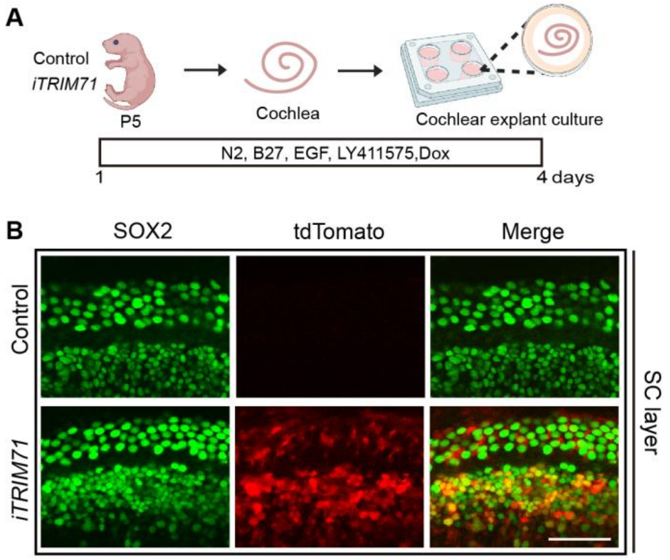


**Fig. S2. TRIM71 transgene expression in cochlear SCs** (*A*) Experimental scheme. To induce TRIM71 transgene expression in cochlear explant culture, Dox was added to the culture medium for the entire culture. Created in BioRender. (*B*) Confocal images of the middle turn of control and TRIM71-overexpressing cochlear explants in the SC layer, immunostained for SOX2 (green). tdTomato was used to monitor TRIM71 expression. Scale bar, 40 μm.

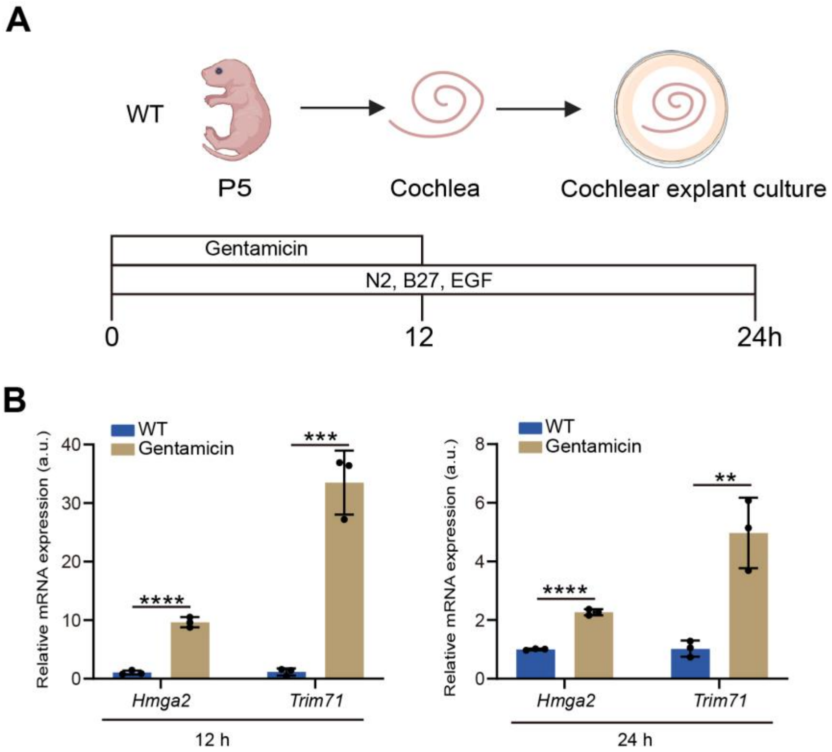


**Fig. S3. HC damage activates *Trim71* expression in cochlear explant culture.** (*A*) Experimental scheme. Cochlear explants were established with P5 mice, and HCs were damaged with gentamicin for 12h. Cochlear epithelia were harvested 12 and 24 hours after the initial gentamicin addition. Created in BioRender. (*B*) RT-qPCR detection of *Hmga2* and *Trim71*. n=3, two independent experiments. Two-tailed unpaired Student's t test was used to calculate *P* values. ***P<0.01, ***P<0.001* and ******P<0.0001.*

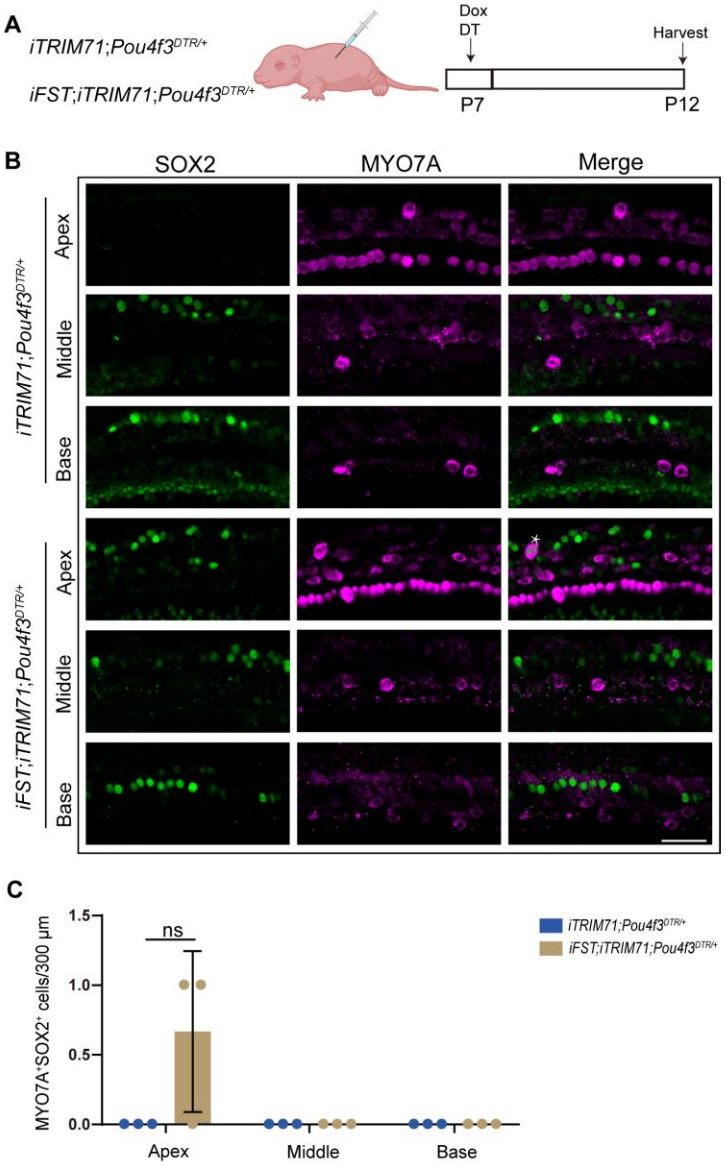


**Fig. S4. Overexpression of TRIM71 and FST fails to boost HC regeneration in stage P7 mice.** (*A*) Experimental scheme. To ablate HCs, mice received DT (6.25 ng/g) at stage P7. To induce FST and/or TRIM71 transgene expression in neonatal mice, mice received Dox (7.5 mg/kg) at stage P7 and harvested tissue at P12. Created in BioRender. (*B*) Confocal images of HC-damaged cochlea isolated from *iTRIM71;Pou4f3*$^{DTR/+}$ and *iFST;iTRIM71;Pou4f3*$^{DTR/+}$ mice at stage P12 stained for MYO7A (magenta) and SOX2 (green). White asterisks indicate SOX2$^{+}$MYO7A$^{+}$ cells. Scale bar, 40 μm. (C) Quantification of new HCs (SOX2$^{+}$MYO7A$^{+}$) formed in the cochlear apex, middle, and base. n=3, two independent experiments. Two-tailed, unpaired Student's *t* test was used to calculate *P* values. n.s., not significant.

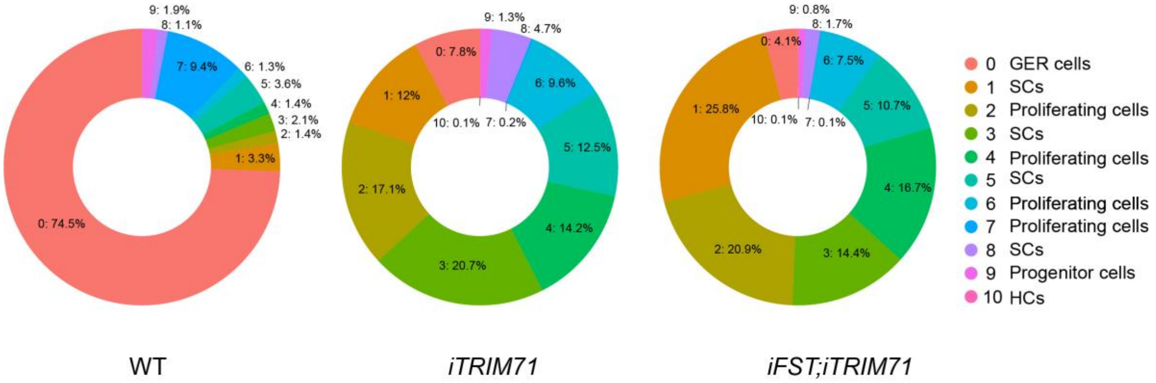


**Fig. S5. Cellular cluster compositions for control, TRIM71 and FST+TRIM71-overexpressing cochlear organoids at 10 days of expansion.** The pie charts show the percentage of cells among different clusters in control, TRIM71, and FST+TRIM71-overexpressing organoids. The cell cluster 0 represents GER cells; 1, 3, 5, 8 represent SCs; 2, 4, 6, 7 represent proliferating cells; 9 represents progenitor cells; 10 represents HCs.

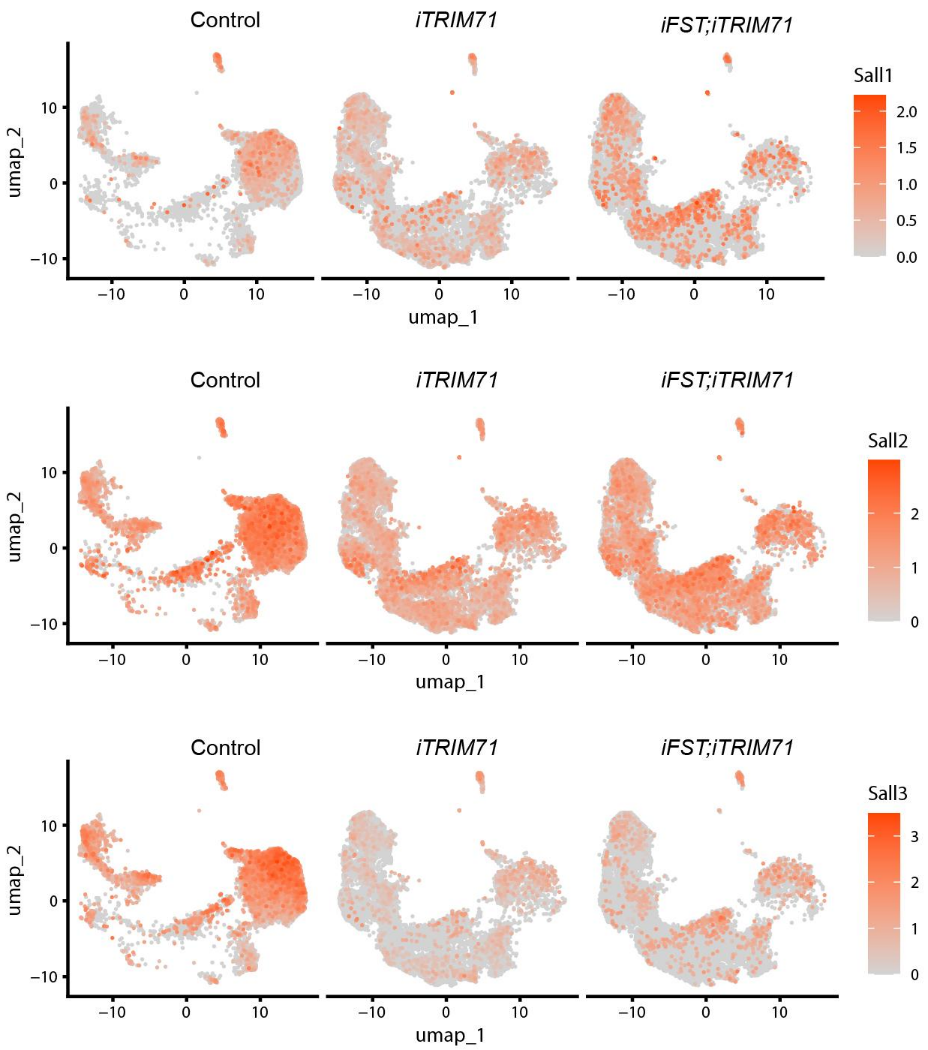


**Fig. S6. Feature plots for *Sall1*, *Sall2,* and *Sall3* expression in control*, iTRIM71,* and i*FST*;i*TRIM71*-overexpressing cochlear organoids at 10 days of expansion.** Organoid cultures were established with cochlear epithelial cells from stage P5 control, *iTRIM71*, and *iFST;iTRIM71* transgenic mice.

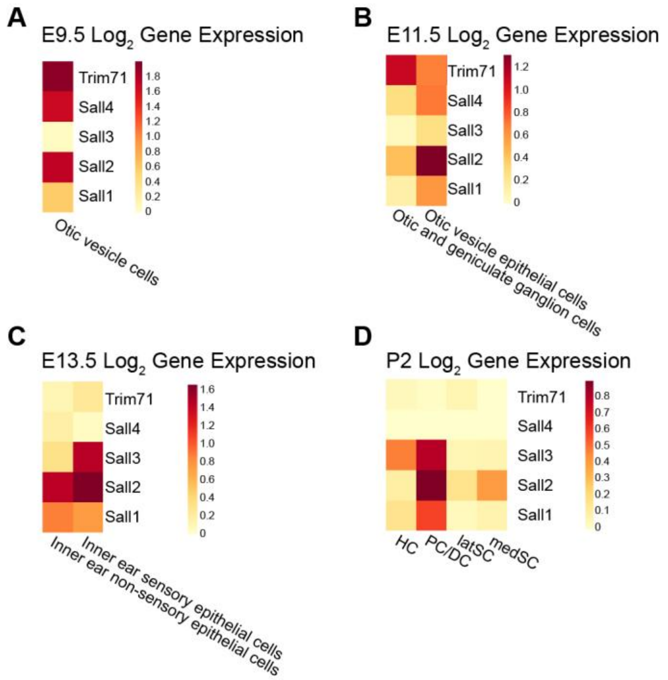


**Fig. S7. Expression of *Trim71* and *Sall1-4* in the developing inner ear and early postnatal cochlea.** (*A*) Heatmap shows the relative expression level of *Trim71*, *Sall1*, *Sall2*, *Sall3* and *Sall4* in the otic vesicle cells at E9.5 (9). (*B*) Heatmap shows the expression of *Trim71*, *Sall1*, *Sall2*, *Sall3,* and *Sall4* in the otic vesicle epithelial cells, otic and geniculate ganglion cells at E11.5 (9). (*C*) Heatmap shows the expression of *Trim71*, *Sall1*, *Sall2*, *Sall3* and *Sall4* in the inner ear sensory and non-sensory epithelial cells at E13.5 (9). (*D*) Heatmap shows the expression of *Trim71*, *Sall1*, *Sall2*, *Sall3,* and *Sall4* in HC, Pillar cells/Deiters' cells (PC/DC), lateral sulcus cells (latSC), and medial sulcus cells (medSC) at P2(10).

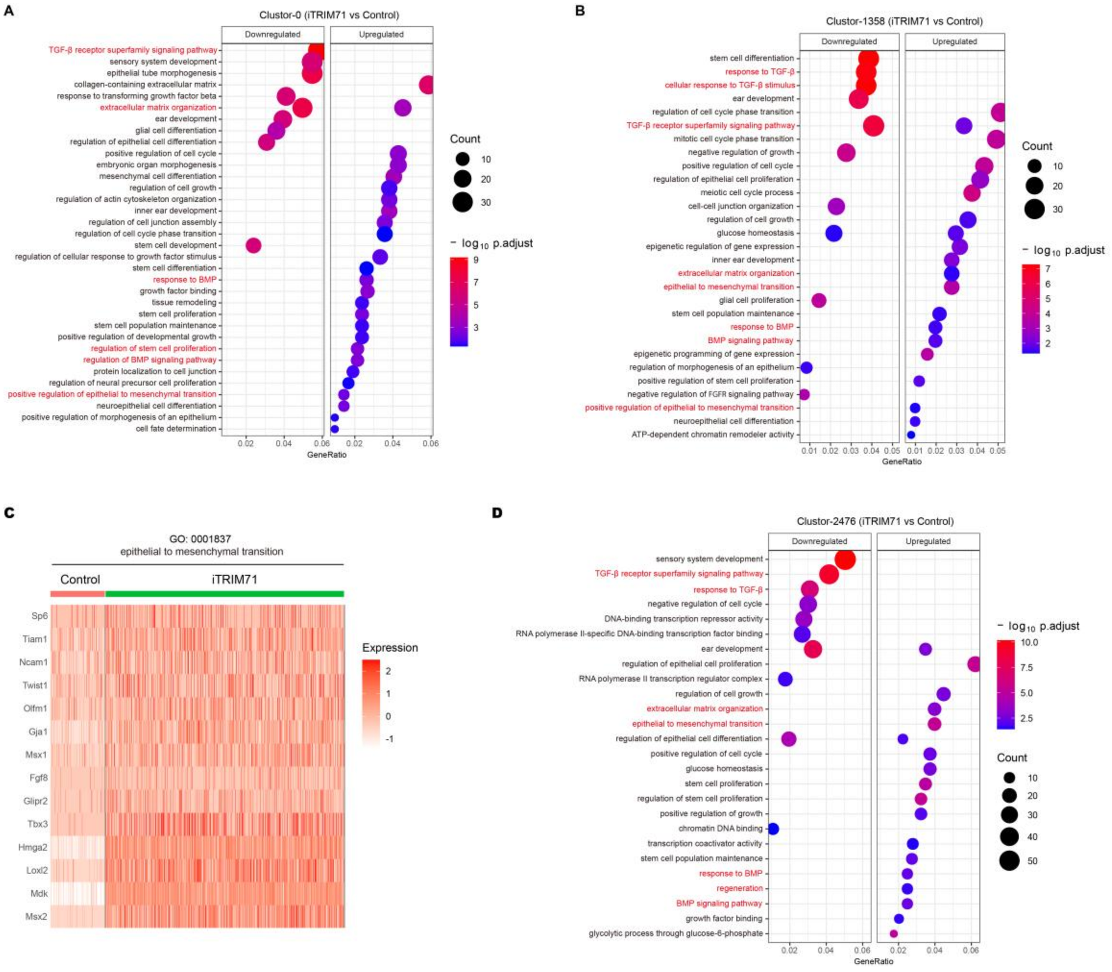


**Fig. S8. TRIM71 overexpression upregulates genes involved in epithelial-to-mesenchymal transition in cochlear organoids.** (*A-D*) Gene Ontology analysis of genes that are significantly changed after TRIM71 overexpression in cluster-0 (*A*), clusters-1,3,5,8 (*B*), and clusters-2,4,7,6 (*D*). (*C*) Heatmap displaying differentially expressed genes (DEGs) in clusters-1,3,5,8 associated with GO:0001837: epithelial-to-mesenchymal transition. The GO terms with adjusted *P* values less than 0.05 were considered significantly enriched among the query genes.

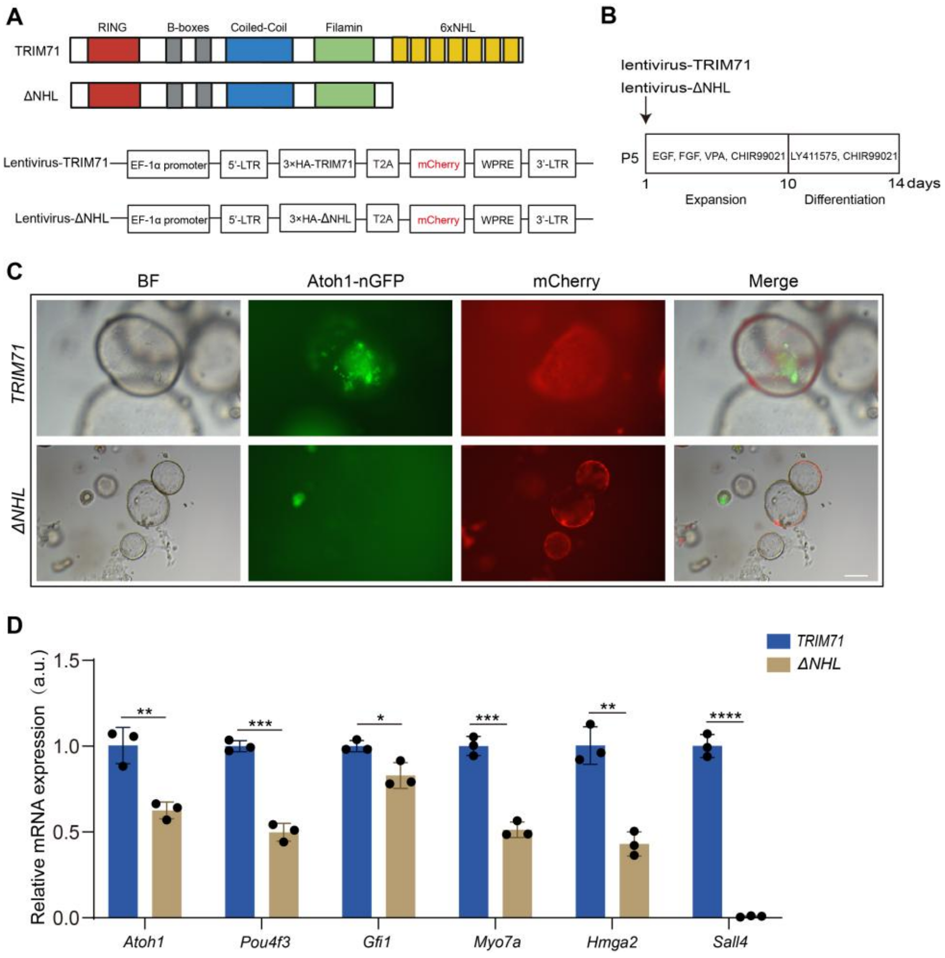


**Fig. S9. The RNA-binding domain of TRIM71 is required for Sall4 activation.** (*A*) Structure of the lentivirus vectors for expressing TRIM71 and ΔNHL. (B) Experimental scheme. Cochlear epithelial cells from P5 *Atoh1-nGFP* tg mice were used to establish organoid cultures, and the cells were infected with lentivirus at the start of culture. (*C*) Low-power bright-field, green (Atoh1-nGFP), and red (mCherry) fluorescent images of organoid culture after TRIM71 and ΔNHL overexpression at 14 days of culture. Scale bar, 200 µm. (*D*) RT-qPCR analysis of the mRNA expression of *Atoh1*, *Pou4f3*, *Gfi1*, *Myo7a*, *Hmga2,* and *Sall4* in TRIM71 and ΔNHL-overexpressing cochlear organoids. n=3, two independent experiments. Two-tailed, unpaired Student's *t* test was used to calculate *P* values. **P<0.05*, ***P<0.01*, ****P<0.001* and *****P<0.0001*.

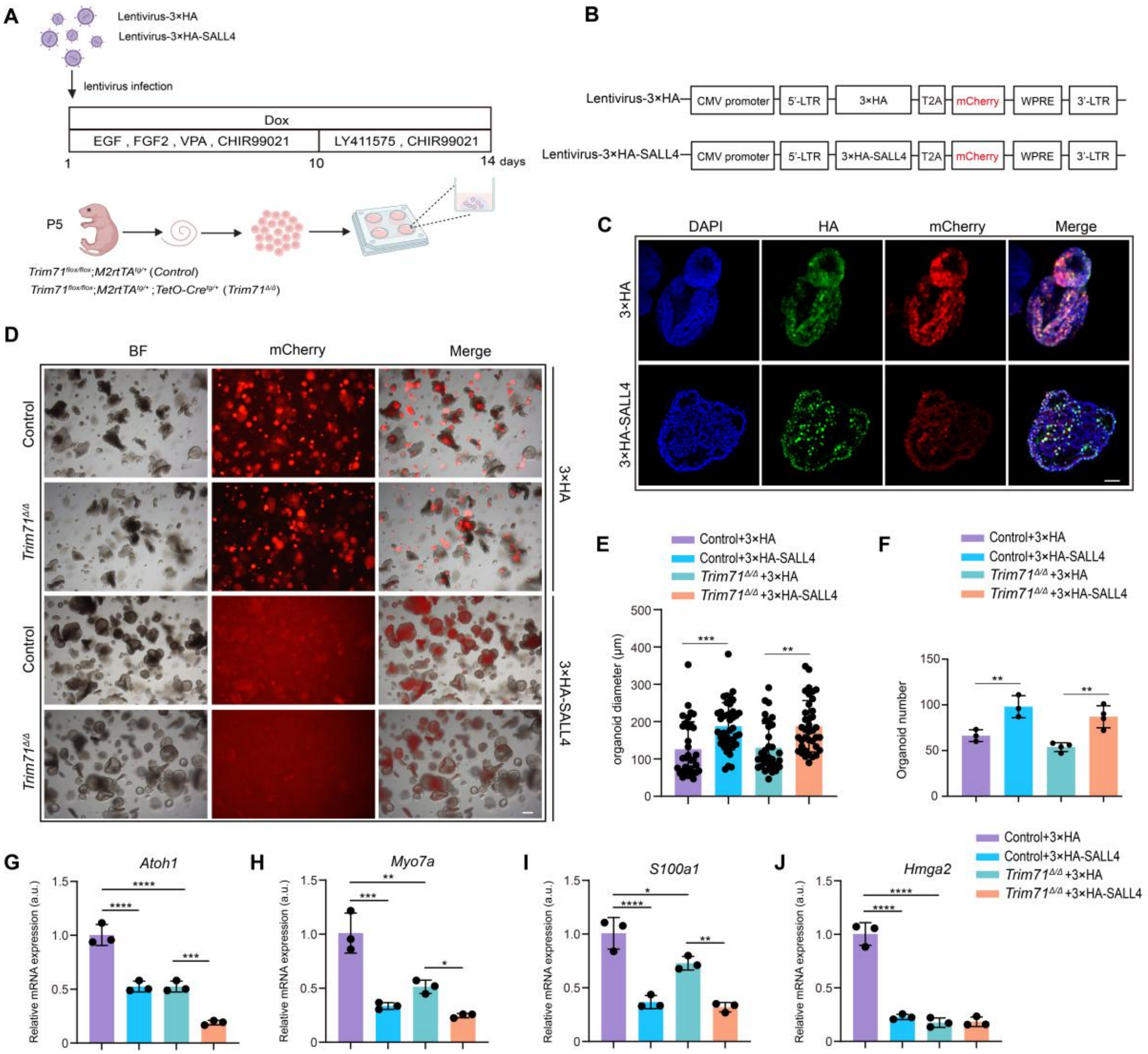


**Fig. S10. SALL4 overexpression promotes cochlear organoid proliferation but inhibits HC formation.** (*A*) Experimental scheme. P5 control (*Trim71*$^{fl/fl}$;*M2rtTA*$^{tg/+}$) and *Trim71*$^{\Delta/\Delta}$ (*Trim71*$^{fl/fl}$;*M2rtTA*$^{tg/+}$;*TetO-Cre*$^{tg/+}$) cochlear epithelial cells were used to establish organoid cultures. Cells were infected with control (HA-tag only) or HA-SALL4-expressing lentivirus at the start of culture. Created in BioRender. (*B*) Structure of lentiviral vectors for SALL4 expression (3×HA-SALL4) and control (3×HA). Both control and SALL4-expressing vectors express mCherry to label infected cells. (*C*) Confocal images of HA-tag in 3×HA and 3×HA-SALL4-overexpressing organoids. Scale bar, 40 μm. (*D*) Brightfield images of organoids after overexpression of 3HA and 3HA-SALL4 in control and *Trim71*$^{\Delta/\Delta}$ mice. mCherry was used to monitor lentivirus infection. Scale bar, 200 μm. (*E*) Organoid diameters in (*D*). (*F*) Organoid number in (*D*). (*G-J*) RT-qPCR analyzing the mRNA expression in control and *Trim71*$^{\Delta/\Delta}$ mice after overexpressing 3×HA and 3×HA-SALL4. n=3, two independent experiments. Two-tailed, unpaired Student's *t* test was used for *E* and *F*. One-way ANOVA and Tukey's multiple comparison test were performed for *G-J*. *$*P<0.05$, $**P<0.01$, $***P<0.001$, $****P<0.0001$.*

### Tables S1 to S4

#### Table 1. Genotyping primers

| Mouse line | Genotyping primers | Product size |
| --- | --- | --- |
| *Atoh1-GFP* | EGFP1: CGA AGG CTA CGT CCA GGA GCG CAC<br>EGFP2: GCA CGG GGC CGT CGC CGA TGG GGG TGT | TG= 300 bp |
| *R26-rtTA*M2* | MTR: GCG AAG AGT TTG TCC TCA ACC<br>F: AAA GTC GCT CTG AGT TGT TAT<br>WTR: GGA GCG GGA GAA ATG GAT ATG | WT=650 bp<br>MT=340 bp |
| *iFST* | YA88: TTG CCT CCT GCT GCT GCT GC<br>YA123: TTT TTC CCA GGT CCA CAG TCC ACG | TG=247 bp |
| *Pou4f3* $^{DTR/+}$ | Common: AAG AAG CAG GTG GGG GAG AG<br>Wild type Reverse: ATT GTT CTG GGC GAC ATG A<br>Mutant Reverse: CAG AAA GAG CTT CAG CAC CAC | Mutant: 290 bp<br>WT: 351 bp |
| *iTRIM71* | F: TGC TGT CTC TTT ATG AGG AGT TGT<br>R: AGA ATG ACA CCT ACT CAG ACA ATG C | TG=586 bp |
| *Trim71-floxed* | Trim71-F-WT: GAA AGG AGG CTA GCC AAA GG<br>Trim71-R-KO: ATG CTG TAC GGT AGG AGT CTT CC | FL=350 bp<br>WT=250 bp |
| *TetO-Cre* | F: GCC TGC ATT ACC GGT CGA TGC AAC GA<br>R: GTG GCA GAT GGC GCG GCA ACA CCA TT | TG=700 bp |

#### Table 2. RT-qPCR primers

| Gene | Forward Primer | Reveres Primer |
| --- | --- | --- |
| *Atoh1* | ATG CAC GGG CTG AAC CA | TCG TTG TTG AAG GAC GGG ATA |
| *Pou4f3* | GCA CCA TCT GCA GGT TCG A | CCG GCT TGA GAG CGA TCA T |
| *Myo7a* | CCC CCT CTG AGA AGT TCG TTA A | TGT GTC CGA GTT CCG TTG AC |
| *Hmga2* | CAG AAG AAA GCA GAG ACC ATT | TTG TTG TGG CCA TTT CCT AGG T |
| *Trim71* | ATC GGG AGT GTG AGC TGT TG | GGC GTG AAC ATA ATG CGG TC |
| *S100a1* | TGG ATG TCC AGA AGG ATG CA | CCG TTT TCA TCC AGT TCC TTC A |
| *Gfi1* | AGG AAC GCA GCT TTG ACT GT | TGA GAT CCA CCT TCC TCT GG |
| *Sall4* | CAA GTC ACC AGG GCT CTT CC | TTC ACT GGA GCA CCG GGT T |
| *Scx* | CTT CAC TGC GCT GCG CAC ACT CAT CC | GCT CTC CGT GAC TCT TCA GTG GCA TCC |
| *Mmp14* | CTG CCA TTG CCG CCA TGC AAA A | TGG CGT GGC ACT CTC CCA TAC T |
| *Twist1* | GGA CAA GCT GAG CAA GAT TCA | CGG AGA AGG CGT AGC TGA G |
| *Rpl19* | GGT CTG GTT GGA TCC CAA | TGC CCG GGA ATG GAC AGT CA |

**Table 3. Antibodies**

| Reagent type | Designation | Source | Identifiers | Additional information |
|---|---|---|---|---|
| antibody | Myosin VIIa rabbit polyclonal | Proteus Biosciences | Cat.# 25-6790 | 1:500 dilution |
| antibody | SOX2 goat polyclonal | Santa Cruz | Cat.# sc-17320 | 1:500 dilution |
| antibody | HMGA2 rabbit monoclonal | Cell Signaling Technology | Cat.# 8179S | 1:1000 dilution |
| antibody | HA-tag Mouse Monoclonal | Vazyme | Cat.# RA1004 | 1:500 dilution |
| antibody | donkey anti-rabbit IgG (H+L) Alexa Fluor 647 | Thermo Fisher | Cat.# A31573 | 1:1000 dilution |
| antibody | donkey anti-rabbit IgG (H+L) Alexa Fluor 555 | Thermo Fisher | Cat.# A31572 | 1:1000 dilution |
| antibody | donkey anti-Goat IgG (H+L) Alexa Fluor 405 | Thermo Fisher | Cat.# A48259 | 1:1000 dilution |
| Nuclear stain | DAPI | Beyotime | Cat.# C1002 | 1:1000 dilution |

**Table 4. Sequences of siRNA for Sall4**

| siRNA | Sense sequence (5'-3') | Antisense sequence (5'-3') |
|---|---|---|
| *NC* | UUC UCC GAC AGU GUC ACG U(dT)(dT) | ACG UGA CAC UGU CGG AGA A(dT)(dT) |
| *Sall4-1#* | CAC UCA CAU CUG CAA CAA A(dT)(dT) | UUU GUU GCA GAU GUG AGU G(dT)(dT) |
| *Sall4-2#* | CCU CGG UGU UAG AUG UCA A(dT)(dT) | UUG ACA UCU AAC ACC GAG G(dT)(dT) |
| *Sall4-3#* | GCU AUU UAC CCA AAG GCA A(dT)(dT) | UUG CCU UUG GGU AAA UAG C(dT)(dT) |

### Datasets S1

Datasets S1. (separate file)

Description: HC and progenitor cell-specific genes list.
